# Charged Molecular Glue Discovery Enabled by Targeted Degron Display

**DOI:** 10.1101/2024.09.24.614843

**Authors:** Zhe Zhuang, Woong Sub Byun, Jakub Chrustowicz, Zuzanna Kozicka, Veronica L. Li, Dinah M. Abeja, Katherine A. Donovan, Sara Sepic, Inchul You, Mikołaj Słabicki, Eric S. Fischer, Stephen M. Hinshaw, Benjamin L. Ebert, Brenda A. Schulman, Nathanael S. Gray

## Abstract

Small molecules that induce protein interactions hold tremendous potential as new medicines, as probes for molecular pathways, and as tools for agriculture. Explosive growth of targeted protein degradation (TPD) drug development has spurred renewed interest in proximity-inducing molecules and especially Molecular Glue Degraders (MGDs). These compounds catalyze destruction of disease-causing proteins by reshaping protein surfaces and promoting cooperative binding between ubiquitylating enzymes and target proteins. MGD discovery for pre-defined targets is a major challenge in contemporary drug discovery. The field is limited by a lack of approaches that can exploit charged ligand-binding pockets, thus excluding a major fraction of ubiquitin ligases (E3s) that evolved to recognize exceedingly common acidic and basic degrons. Here we solve these important chemical challenges through “chemocentric” MGD discovery of **ZZ1**, a BET-family protein degrader and a prodrug of a negatively **<u>c</u>**harged **<u>glue</u>** (c-Glue). **ZZ1** activation unmasks a sulfinic acid moiety that binds the modular GID/CTLH ubiquitin ligase complex via a basic pocket in its YPEL5 subunit. YPEL5 is a CRBN structural homolog and an essential non-Cullin ubiquitin ligase cofactor expressed in cancers of the bone marrow. These findings demonstrate a previously unrecognized capacity of YPEL5 to recruit GID/CTLH substrates, and they provide a powerful strategy to discover c-Glues that induce proximity to ubiquitin ligases with similarly desirable properties.

## Main

Discovery of small molecules that induce binding between two or more target proteins is an important challenge with major implications for medicine, agriculture, and basic biology^1-5^. Such molecules are valuable because they can produce unexpected pharmacology by redirecting enzymatic activities to new substrates. Molecular glue degraders (MGDs) embody this concept. These small molecules remodel protein surfaces to promote binding between ubiquitylating enzymes and so-called “neosubstrates,” ultimately resulting in targeted protein degradation (TPD)^2^. MGDs have superior drug-like properties, induce TPD at low compound concentrations by acting catalytically, do not require pre-existing ligands for the two recruited proteins, and can recruit partners without drug binding pockets^1,6,7^.

Discovery of new MGDs that bind previously undrugged ubiquitylating enzymes is a prominent challenge in contemporary chemical biology^8-11^. Chemical elaboration of known MGDs to refocus their activities on distinct neosubstrate targets is a proven drug discovery strategy and motivates this search^12-14^. This is an especially important challenge in the MGD space, because the ubiquitin ligase-MGD “neo-surface” (rather than the compound alone) determines which neosubstrate surfaces bind and how these proteins are presented to ubiquitylation active sites^15^. Further, different ligase-MGD pairs may bind distinct neosubstrate surfaces on the same target protein. A consequence of this is that each ligase-MGD pair engages an idiosyncratic and unpredictable assortment of neosubstrate proteins and therefore offers unique off-target and, ultimately, genetic/epigenetic drug resistance profiles. Tissue-restricted expression of ubiquitin ligase components adds additional opportunities for MGD-neosubstrate selectivity. Despite these possibilities, most MGDs bind ubiquitylating enzymes in the cullin-RING ligase family that are broadly expressed and not required for cell proliferation. The dual consequences are drug resistance by ligase downregulation and poor therapeutic indices due to systemic TPD activity^16^.

Current approaches to MGD discovery exclude a major fraction of all E3 ligases: those that bind charged clients. In fact, many of the first E3 ligase families discovered degrade cellular substrates with positive or negative charges. Early examples include N-end rule degrons^17-19^ and phosphodegrons^20,21^. Relatively more recently identified E3 ligases recognize C-terminal degrons^22-24^ or clusters of acidic or basic residues^25,26^. Participation of these E3 ligase systems in cell cycle signaling, protein quality control, metabolism, and cellular differentiation underlines their physiologic prevalence and importance.^17-27^

Substrate-mimicking MGDs bearing complementary charges are needed to address the large group of E3 ligase systems dependent on electrostatically driven substrate binding. Such molecules have not emerged from previous MGD discovery campaigns. This is largely due to difficulties in developing cell permeant charged small molecules, which is a recurring and general challenge in medicinal chemistry. MGD discovery commonly entails expansion of the neosubstrate scopes of known E3 ligase-MGD pairs^12-14^. In a few cases, cryptic target depletion by potent small molecule inhibitors has led to MGD discovery^28-31^. Both strategies are limited to degrader discovery, and neither are generalizable to glue molecules with other pharmacologies, nor for developing new molecular glues for pre-defined target proteins. We and others have shown that subtle chemical modifications can endow gain-of-function MGD properties upon existing chemical ligands^32,33^. Molecules discovered by this “chemocentric” strategy primarily target Cullin-RING family E3 ligases. High resolution structures of ligase-small molecule-neosubstrate ternary complexes are essential for targeted MGD diversification. Except in rare cases^30^, molecular structures have not been reported for most MGDs discovered via this strategy.

Here, we used a target-focused reactive chemical screening strategy, coupled with counter-screening, to identify a protein degradation mechanism that has not yet been exploited for TPD. The resultant **ZZ1** molecule and the rationally improved **ZZ2** molecule are metabolically activated, target the previously undrugged YPEL5-GID/CTLH E3 complex, and are **<u>c</u>**harged **<u>glue</u>**(c-Glue) small molecules promoting electrostatically-induced proximity.

### Results: Discovery of a YPEL5-GID/CTLH-dependent BRD4 degrader

To identify MGDs that exploit novel E3 ligases, we employed a degron display approach whereby we functionalized a known high-affinity ligand of the target protein with a library of chemical tags containing covalent warheads and heterocycles. We chose the bromodomain and extra-terminal domain (BET) family of epigenetic readers as model targets, and we used the BET bromodomain inhibitor JQ1 as a parental substrate-recruiting handle^34^. This BRD4-JQ1 system is an established system for discovery of monovalent glue degraders^30-32^. We detected BRD4 degradation by a HiBiT assay based on CRISPR/Cas9-mediated fusion of an 11-amino acid peptide-tag to BRD4^32^. A relatively simple chemical tag appended to JQ1 yielded a functional degrader that we refer to as ZZ1 (Fig. 1a). ZZ1 features an amide-linked phenyl ring with chloro and sulfonyl fluoride^35^ groups at the *meta* and *para* positions, respectively. ZZ1 induced selective degradation of BET proteins (BRD3 and BRD4) as determined by immunoblotting and quantitative proteome-wide mass spectrometry (Extended Data Fig. 1a, 1b).

**Figure 1:**
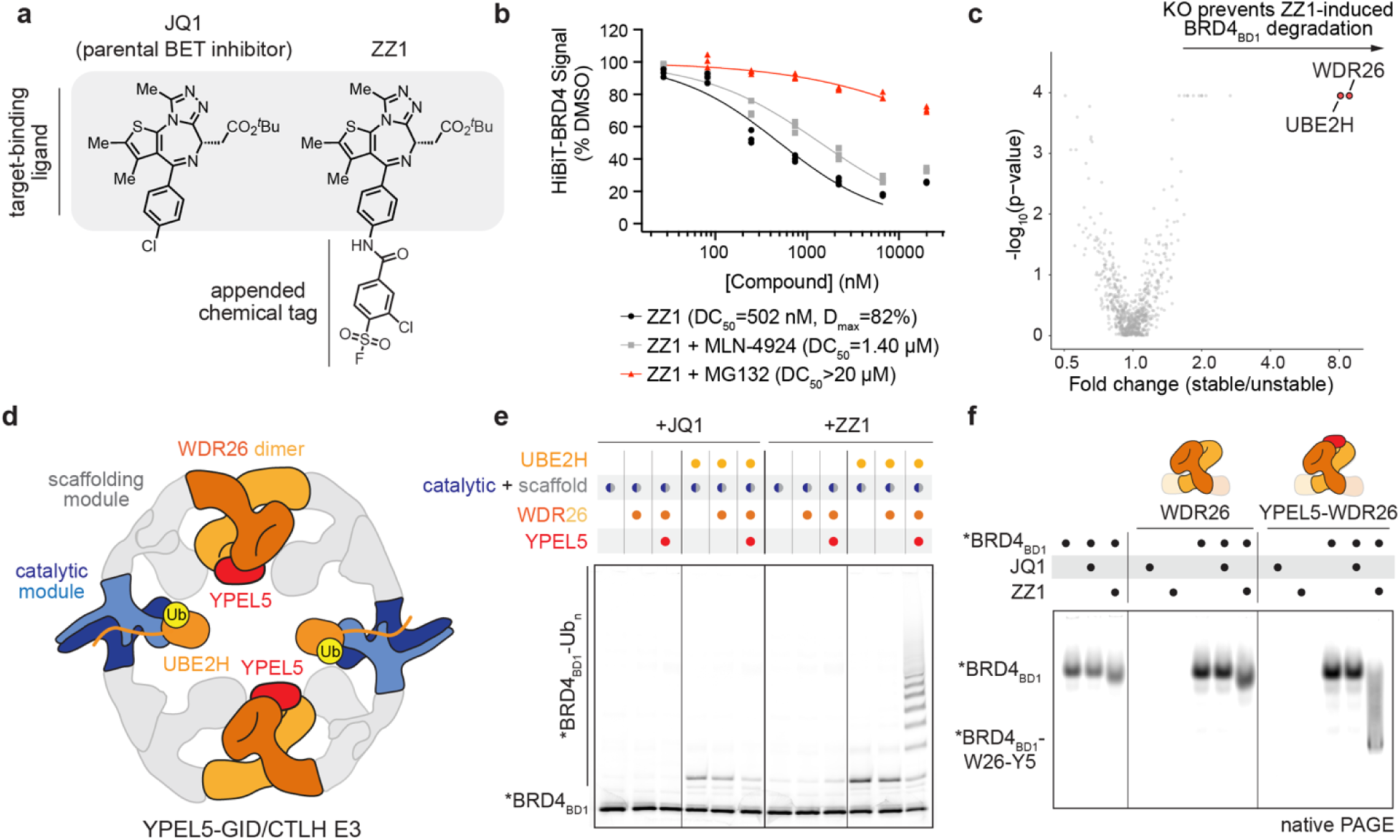
Chemocentric approach yields a molecular glue degrader for a pre-defined target. a) Chemical structures of the parental BET protein family inhibitor (JQ1) and its derivative (ZZ1) featuring an appended chemical tag conferring degrader activity. b) HiBiT-BRD4 assay results for Jurkat cells pre-treated with the indicated inhibitors for 1 h, followed by treatment with ZZ1 for 5 h. c) Ubiquitin-proteasome system (UPS)-focused CRISPR screen for BRD4_BD1_-eGFP stability in K562-Cas9 cells treated with 1 µM ZZ1 for 16 h. d) Cartoon of the UBE2H∼Ub-bound YPEL5-GID/CTLH E3 ligase catalytic assembly highlighting its functional modules. e) *In vitro* ubiquitylation assay of fluorescently labeled *BRD4_BD1_ (asterisk denotes an N-terminal FAM label) determining the minimal catalytic assembly sufficient for ZZ1-dependent activity. The examined GID/CTLH E3 ligases comprised the catalytic core (catalytic and scaffolding modules) alone or assembled with substrate regulatory subunits WDR26 and YPEL5. Reactions were quenched after 45 min. See Extended Data Figure 2 for details of the GID/CTLH E3 ligase architecture and assays with a full suite of GID/CTLH E3 assemblies. f) Native gel mobility shift assay probing ZZ1-induced *BRD4_BD1_ engagement by the WDR26 dimer or WDR26-YPEL5. Transparent regions in the cartoon represent truncated WDR26 domains. The truncations prevent higher-order oligomerization).

Evidence supporting a novel mechanism of action for ZZ1 surfaced during further examination of BRD4 destabilization. First, ZZ1-induced BRD4 degradation in the HiBiT assay was not sensitive to cullin-RING ubiquitin ligase (CRL) inhibition (via NEDD8 activating enzyme inhibitor MLN-4924), while proteasome inhibition prevented degradation (Fig. 1b). These data suggest involvement of a novel non-CRL E3 ligase. Second, ZZ1 induced selective degradation of the bromodomain. We used a fluorescent reporter assay to identify the region of BRD4 responsible for ZZ1-induced degradation (Extended Data Fig. 1c)^30^. Despite the similar affinity of JQ1 to both BRD4 bromodomains (BDs), only reporters containing bromodomain 1 (BRD4_BD1_ as well as the tandem construct containing both BD1 and BD2, BRD4_BD1+BD2_) were degraded. Thus, ZZ1 harnesses a distinct BD compared to most JQ1-based degraders^30-32,36^, implying involvement of a previously undrugged E3 ligase.

To identify ubiquitylation factors required for ZZ1-induced BRD4_BD1_ degradation, we performed a fluorescence-activated cell sorting CRISPR screen, testing 713 genes involved in the ubiquitin-proteasome system (UPS)^28^ (Fig. 1c). The screen identified key components of the GID/CTLH E3 ligase, including its cognate E2 enzyme, UBE2H^37,38^, and its subunit, WDR26^39,40^ (Fig. 1d, Extended Data Fig. 2a). To determine whether the GID/CTLH E3 can ubiquitylate BRD4, we reconstituted the ZZ1-mediated activity *in vitro* using purified proteins. GID/CTLH is not a singular E3 ligase but a collection of assemblies with a common catalytic core and varying auxiliary subunits that control substrate selection^39,41^ (Extended Data Fig. 2a, 2b). Testing of all known GID/CTLH auxiliary subunits enabled identification of a minimal assembly proficient to catalyze UBE2H-mediated BRD4_BD1_ ubiquitylation. This complex features WDR26 and its binding partner YPEL5 (Fig. 1e, Extended Data Fig. 2c). Accordingly, Cas9-mediated knockout of either WDR26 or YPEL5 prevented ZZ1-induced BRD4 degradation (Extended Data Fig. 2d, 2e). Consistent with this finding, ZZ1 is especially potent in the Ewing’s sarcoma TC-71 cell line, which has elevated YPEL5 expression^42^ (Extended Data Fig. 2f). A co-immunoprecipitation experiment using BRD4 as a bait confirmed the presence of the YPEL5-WDR26 regulatory module in the GID/CTLH E3 assembly that engages BRD4 upon cellular treatment with ZZ1 (Extended Data Fig. 2g). Thus, the GID/CTLH complex containing both WDR26 and YPEL5 is necessary and sufficient for ZZ1-mediated BRD4 degradation.

Biochemical and genetic dependence on YPEL5 raised two questions. First, is YPEL5 a substrate receptor of the GID/CTLH E3? The currently only known endogenous function of YPEL5 is to bind WDR26 and prevent it from recruiting its native substrate^43^ (Extended Data Fig. 2h, 2i). In contrast, YPEL5 appears to be responsible for degrader-induced recruitment of the neosubstrate, which is supported by ZZ1-dependent co-migration of BRD4_BD1_ in a non-denaturing gel shift experiment (Fig. 1f). Second, YPEL5 shows striking structural homology to the thalidomide-binding domain of CRBN (CRBN_CTD_), raising a question whether BRD4-ZZ1 binding occurs via a similar structural mechanism.

### ZZ1 glues BRD4_BD1_ to the CRBN-like domain of YPEL5

To understand the mechanism of ZZ1-induced ternary complex formation, we determined a cryo-EM structure of the fully assembled BRD4_BD1_-bound YPEL5-GID/CTLH complex. As reported previously, the WDR26-containing GID/CTLH E3 is a giant twofold symmetric oval with an external diameter of 300 Å in its longest aspect and a hollow center that accommodates substrates^39,40,43^. The overall map, resolved to ∼12-Å, shows two centrally located BRD4_BD1_ molecules, each interacting with twofold symmetry-related YPEL5-WDR26 modules (Fig. 2a, Extended Data Table 1). This binding mode positions BRD4_BD1_ adjacent to the heterodimeric RING domains of the catalytic GID/CTLH subunits, thus explaining the UBE2H-dependent ubiquitylation. By focused refinement and signal subtraction with a mask over the aligned YPEL5-WDR26 modules bound to BRD4_BD1_-ZZ1, we determine a subcomplex structure resolved to 3.4 Å. This enabled unambiguous modeling of the MGD-driven ternary complex (Extended Data Table 1, Extended Data Fig. 3). The modeled interface showed three remarkable features.

**Figure 2:**
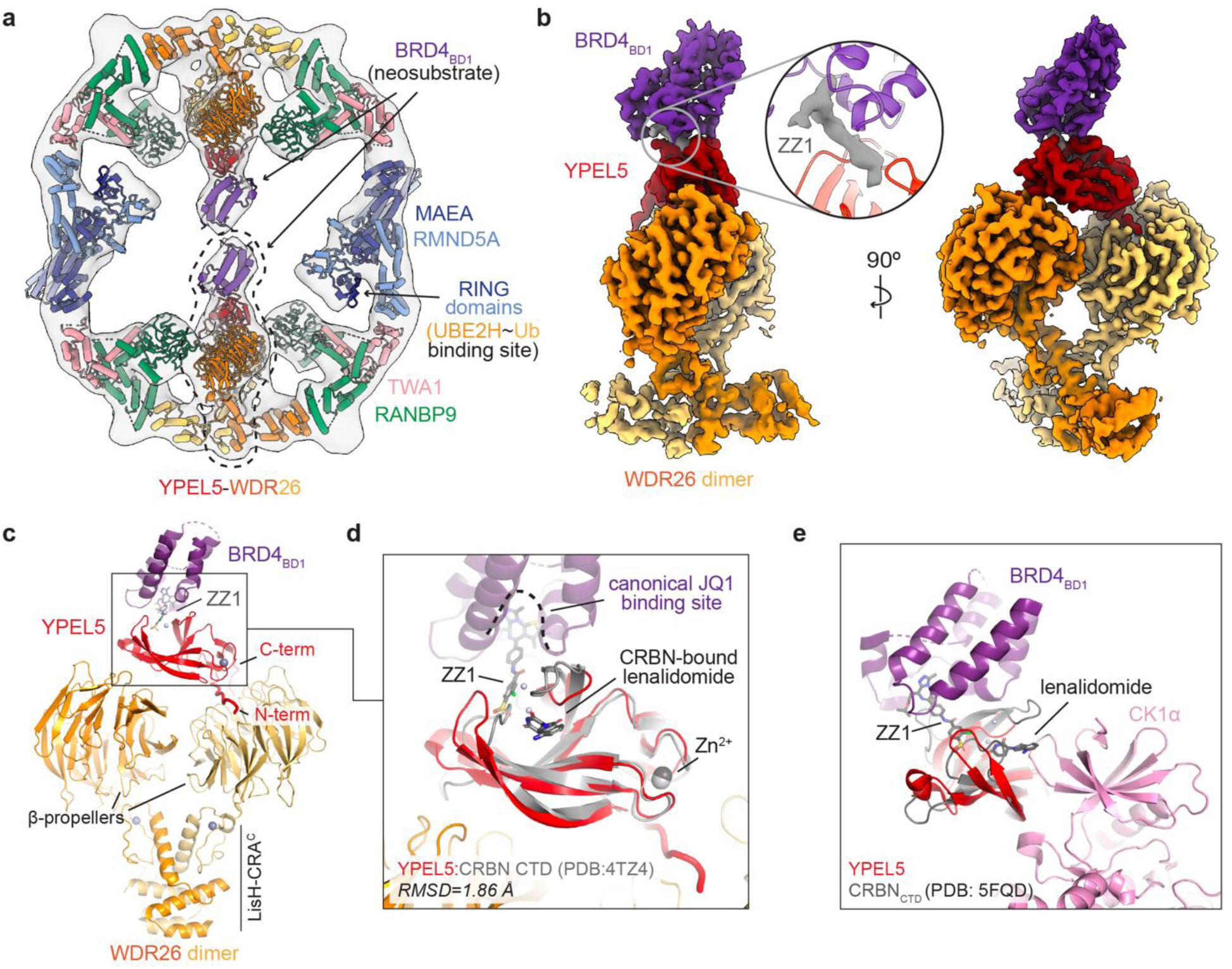
YPEL5 is a ZZ1-induced substrate receptor for BRD4 targeting. a) Cryo-EM map of the neosubstrate recognition complex (YPEL5-GID/CTLH E3-ZZ1-BRD4_BD1_) resolved to 12 Å and fit with prior structures (extracted from PDB: 7NSC^39^, 8PJN^38^, 8QBN^43^, 3MXF^34^) and AlphaFold models^66^ of the constituent GID/CTLH modules. b) Map of the YPEL5-WDR26 module-ZZ1-BRD4_BD1_ ternary complex resolved to 3.4 Å and sharpened with DeepEMhancer^67^. Close-up highlights additional electron density at the interface of YPEL5 and BRD4_BD1_ corresponding to the ZZ1 degrader. c) Atomic model of the ZZ1-induced ternary complex depicting the overall YPEL5-WDR26 receptor module architecture and its mode of BRD4_BD1_ engagement. d) Close-up of YPEL5 in complex with ZZ1-BRD4_BD1_ overlayed with lenalidomide-bound CRBN_CTD_ (PDB: 4TZ4^68^) illustrating the structural similarity of their ligand-binding domains. Both domains are stabilized by coordination of a zinc atom and contain a central groove that binds ligands. e) YPEL5 and CRBN (PDB: 5FQD^46^) MGD ternary complexes have divergent modes of degrader engagement and neosubstrate positioning.

First, ZZ1 is a MGD that bridges YPEL5 and BRD4_BD1_ (Fig. 2b, 2c). The JQ1 moiety of ZZ1 interacts with its canonical binding site on BRD4_BD1_, and the appended degron protrudes out of the BRD4_BD1_ pocket, creating a neomorphic surface for YPEL5 engagement (Fig. 2d). The exposed chemical moiety inserts into the central β-sheet groove of YPEL5, inducing YPEL5-BRD4_BD1_ protein interactions. Involvement of specific protein-protein contacts explains the preference for BRD4_BD1_ over BRD4_BD2_ in our cellular fluorescent reporter degradation and *in vitro* ubiquitylation assays (Extended Data Fig. 1c, 2c). It also suggested interactions would be conserved specifically with the highly similar BD1 domains of BRD2 and BRD3, which were indeed selectively ubiquitylated (Extended Data Fig. 4a).

Second, ZZ1 binds the ligand-binding pocket in YPEL5 that corresponds to the thalidomide-binding pocket of CRBN (CRBN_CTD_)^43^ (Fig. 2d). YPEL5 and CRBN_CTD_ have high structural similarity (RMSD=1.86 Å for the Lon-N-term protease domain) despite low sequence identity (∼17%). Consistent with the divergent primary sequences, neither the molecular details of degrader engagement (ZZ1 for YPEL5 and thalidomide for CRBN) nor the orientations of the recruited neo-substrates relative to the shared binding groove (Fig. 2e, Extended Data Fig. 4b) are conserved. Consequently, the CRBN-based PROTACs employing the BRD4 ligand JQ1 did not trigger YPEL5-dependent BRD4_BD1_ ubiquitylation (Extended Data Fig. 4c). This is consistent with a lack of cellular competition between the thalidomide analog CC-92480 and ZZ1 for BRD4_BD1_ reporter degradation (Extended Data Fig. 4d). Thus, YPEL5 and CRBN_CTD_, have distinct ligand-binding preferences despite sharing a fold.

### MGD function of ZZ1 requires its metabolism to sulfinic acid

The third feature shown in the cryo-EM structure is that the bound ZZ1 appears to have undergone a chemical transformation that enables YPEL5 binding. The chemical tag of ZZ1 projects into the basic environment of the YPEL5 binding groove (Fig. 3a). This was unexpected, as the sulfonyl fluoride moiety does not carry a complementary negative charge. In addition, we did not observe a chemical link between ZZ1 and any surrounding YPEL5 nucleophilic amino acid side chains, although the density for ZZ1 itself was well-defined (Extended Data Fig. 5a). Accordingly, we found no evidence of a covalent ZZ1-YPEL5 adduct in mass spectrometry experiments carried out with recombinant proteins (Extended Data Fig. 5b). Thus, although ZZ1 features a common electrophilic covalent chemical warhead (sulfonyl fluoride)^35^, its mechanism of action does not involve covalent bond formation.

**Figure 3:**
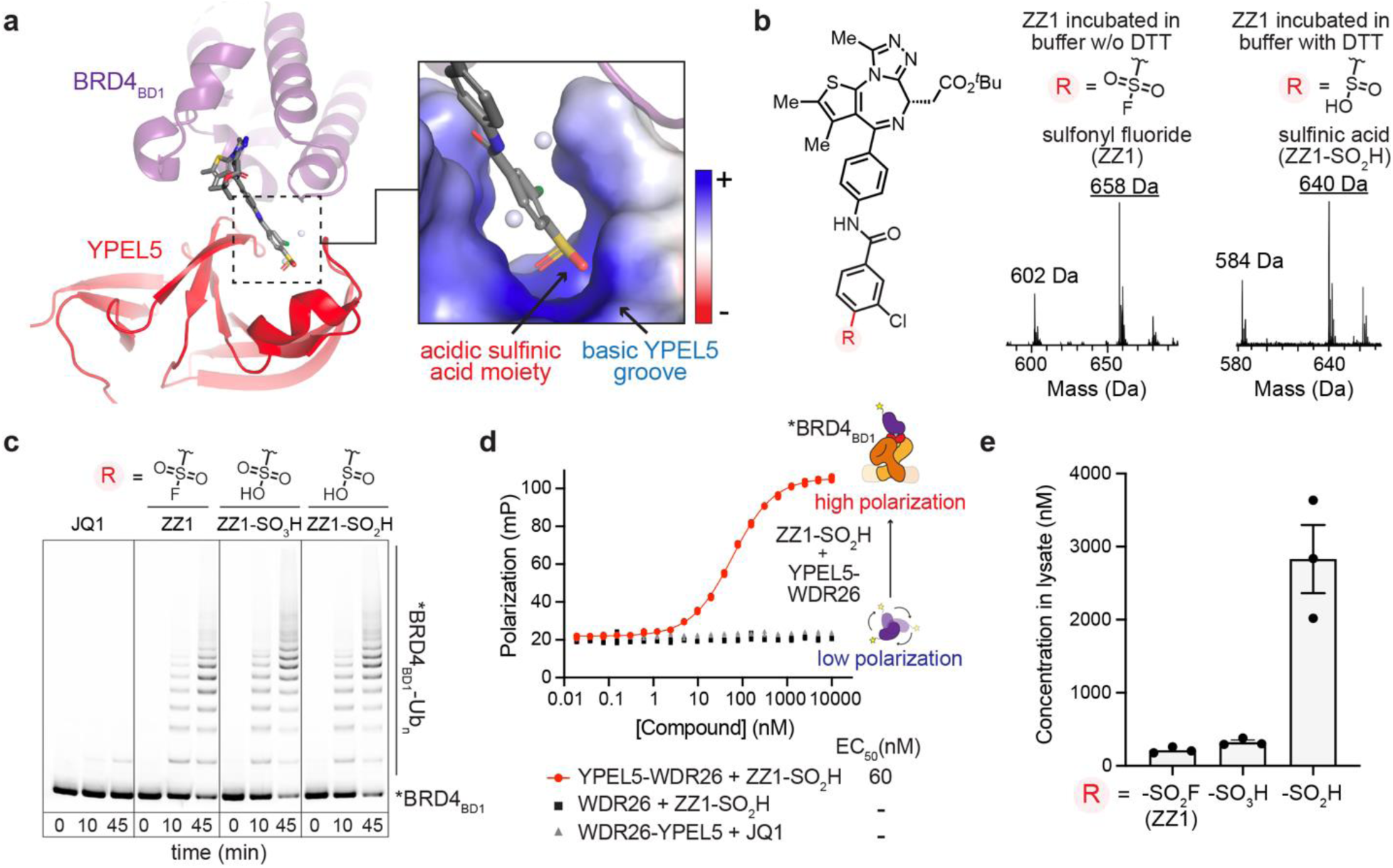
ZZ1 is a metabolically activated pro-drug of a charged molecular glue degrader (c-Glue) a) Structure of the ternary complex illustrating electrostatically-driven interactions between the negatively charged sulfinic acid moiety of ZZ1-SO_2_H and the basic bottom of the YPEL5 binding groove (represented as an electrostatic potential surface). b) Intact mass spectrometry demonstrates conversion of the sulfonyl fluoride moiety of ZZ1 (middle) to sulfinic acid (right) after incubation in a DTT-containing buffer. The position of the transformed group is indicated in the degrader’s chemical structure (left). Peaks corresponding to lower molecular weight species correspond to ZZ1 and ZZ1-SO_2_H derivatives formed upon hydrolysis of their *tert*-butyl ester. c) Stimulation of YPEL5-GID/CTLH E3-dependent *in vitro* ubiquitylation of *BRD4_BD1_. ZZ1, its sulfinic acid derivative, (ZZ1-SO_2_H) and its sulfonic acid derivative (ZZ1-SO_3_H) were tested. d) FP assay quantifying the propensity of ZZ1-SO_2_H to induce the ternary complex formation. Binding of *BRD4_BD1_ to the YPEL5-WDR26 module upon degrader titration results in a dose-dependent increase of fluorescence polarization. Fitting polarization values to the “[agonist] vs. response” model yielded the half-maximal effective concentration (EC_50_). e) HPLC analysis of intracellular levels of ZZ1 and its acidic metabolites. Jurkat cells were treated with 5 µM ZZ1 for 5 h.

We hypothesized that ZZ1 is a prodrug that requires chemical transformation into an acidic moiety for its activity. We tested this idea in several ways. First, mass spectrometry analysis of ZZ1 incubated in the buffer used for biochemical assays indicated the conversion to a sulfinic acid derivative (ZZ1-SO_2_H) (Fig. 3b). Second, both ZZ1-SO_2_H and the related sulfonic acid version (ZZ1-SO_3_H) promoted robust BRD4_BD1_ ubiquitylation *in vitro* (Fig. 3c). Third, we established a fluorescence polarization assay to quantify the propensity of ZZ1 to drive ternary complex formation. The assay measures binding of fluorescently labeled BRD4_BD1_ to YPEL5-WDR26. ZZ1-SO_2_H induces a tight ternary complex between YPEL5-WDR26 module and BRD4_BD1_ with EC_50_ of ∼60 nM (Fig. 3d). Fourth, ZZ1 induced ternary complex formation slowly in biochemical assays when compared with ZZ1-SO_2_H (Extended Data Fig. 5c). Pre-incubating ZZ1 in the reaction buffer alleviated the delay. Fifth, since sulfonyl fluoride is a potent electrophile, we speculated it might readily react with the nucleophilic thiol groups of the reducing agent dithiothreitol (DTT) present in the assay buffer to yield ZZ1-SO_2_H (Extended Data Fig. 5d). Indeed, ZZ1 transformation *in vitro* occurred only in the presence of the reducing agent DTT, with the concentration of DTT correlating with the rate of ZZ1-dependent ternary complex formation (Fig. 3b, Extended Data Fig. 5e). A similar reaction could take place in the cellular environment, potentially mediated by other thiol-containing molecules such as glutathione (GSH) whose involvement is supported by subtle enhancement of ZZ1 potency in GSH pre-treated cells^44,45^ (Extended Data Fig. 5f). Sixth and finally, metabolomic analyses of lysates from ZZ1-treated cells identified ZZ1-SO_2_H as the predominating intracellular species versus the non-converted parental compound (Fig. 3e).

Given the evidence outlined above, we used the converted ZZ1-SO_2_H to model the cryo-EM density and inspected the resulting model to determine the basis of molecular recognition (Fig. 4a, 4b). First, electrostatic contacts between the acidic ZZ1-SO_2_H moiety and basic YPEL5 pocket anchor the complex. Accordingly, increasing the basic character of the pocket by mutating Lys95 to Arg potentiated ZZ1-induced BRD_BD1_ ubiquitylation, and mutating the adjacent Thr63 with a negatively charged Asp attenuated this activity (Fig. 4c). Second, hydrophobic residues within the loops lining the entrance to the YPEL5 groove sandwich the phenyl group of the degron tag. Disrupting these hydrophobic contacts by mutating Leu62, which interacts with both ZZ1 and the surrounding apolar BRD4 residues, nearly completely abolished ZZ1 activity *in vitro* and considerably impaired BRD4 degradation (Fig. 4c, 4d, Extended Data Fig. 6a). Finally, the amide linker between the BRD4 ligand and the appended chemical tag forms a hydrogen bond with YPEL5 Thr36, thereby buttressing YPEL5 engagement. Mutation of YPEL5 at this position weakened ZZ1 potency *in vivo* (Fig. 4d, Extended Data Fig. 6a). Therefore, sulfinic acid recognition is the defining feature of ZZ1-SO_2_H recognition, and secondary interactions with the compound support this contact. Designation of ZZ1 as a c-Glue emphasizes this arrangement.

**Figure 4:**
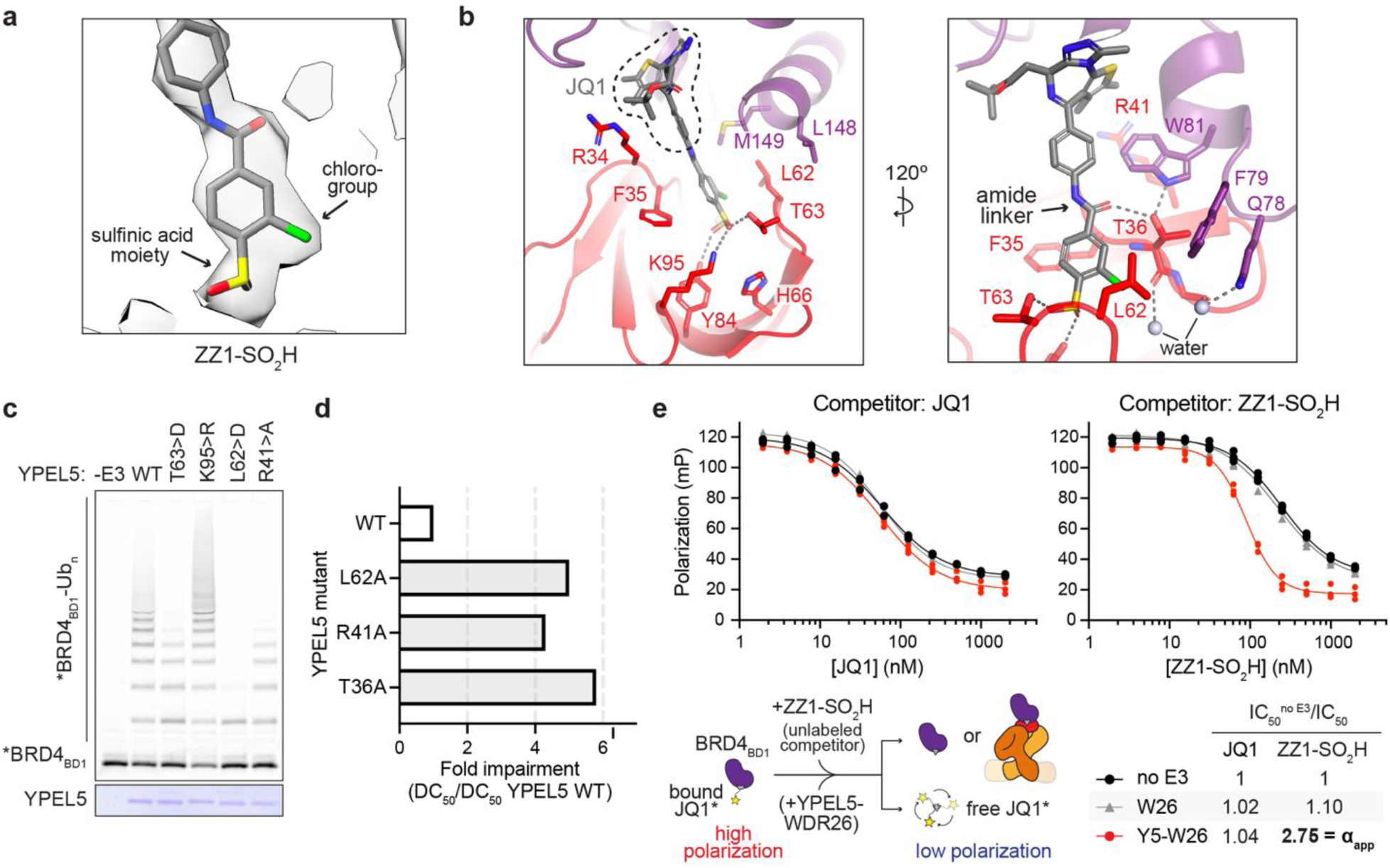
ZZ1-SO_2_H induces cooperative ternary complex formation. a) Close-up of cryo-EM density corresponding to the YPEL5-engaging chemical tag of ZZ1-SO_2_H (compound coordinates shown as sticks). b) Molecular details of the ternary complex interface highlighting the constellation of YPEL5 residues engaging ZZ1-SO_2_H and those involved in direct interactions with BRD4_BD1_. The hydrogen bonds are depicted as gray dashes. c) *In vitro* ubiquitylation assay probing YPEL5 residues shown in (b) involved in: (1) anchoring the sulfinic acid moiety (T63, K95), (2) contacts with the degrader phenyl ring and BRD4 hydrophobic sidechains (L62), and (3) direct interactions with BRD4 (R41). Reactions were quenched after 30 minutes. SDS-PAGE gels were imaged by a fluorescence scan and stained with Coomassie to visually inspect YPEL5 levels (bottom). d) Cellular BRD4 degradation assay testing the structurally visualized binding mode. The impact of YPEL5 mutations on ZZ1 potency is illustrated as the ratio of DC_50_ values between mutant and WT YPEL5-expressing Jurkat cells. Plots used for DC_50_ measurements are presented in Extended Data Figure 6a. e) Competitive FP assay probing cooperativity within the ZZ1-SO_2_H-induced ternary complex. BRD4_BD1_-bound fluorescent JQ1* tracer was displaced by titration of unlabeled competitors reducing fluorescence polarization. The extent of cooperativity was determined by calculating the ratio of IC_50_ values (estimated by fitting polarization values to the “[inhibitor] vs. response” model) in the absence and presence of excess YPEL5-WDR26 (apparent cooperativity factor α_app_).

Akin to other structurally characterized MGDs^12,46^, ZZ1-SO_2_H promotes direct target protein-E3 ligase contacts^47^, burying a total surface area of ∼350 Å^2^. These include both polar interactions at the water-mediated three-way interface of ZZ1’s chloro group, YPEL5, and BRD4, as well as hydrophobic packing of YPEL5 Arg41 against BRD4 Trp81 at the other side of the assembly (Fig. 4b). Mutating YPEL5 Arg41 to Ala attenuated ZZ1 MGD activity *in vitro* and in cells (Fig. 4c, 4d, Extended Data Fig. 6a). To further probe the importance of protein-protein contacts for ZZ1-induced ternary complex, we employed a competitive fluorescence polarization assay (Fig. 4e, Extended Data Fig. 6b). Whereas JQ1 displaced the fluorescent JQ1 probe from BRD4_BD1_ with similar IC_50_ regardless of E3 ligase presence, ZZ1-SO_2_H exhibited a 2.75-fold stronger suppressive effect (corresponding to an apparent cooperativity factor α_app_) in the presence of YPEL5-WDR26 module but not WDR26 alone. The enhanced affinity of a degrader to a target protein in the presence of an E3 ligase indicates the formation of a positively cooperative ternary complex, corroborating the structurally observed binding mode.

### Structure-based optimization yields the superior degrader ZZ2

We hypothesized that introducing a small electronegative group into a vacant basic pocket adjacent to the unsubstituted *ortho*-position of the ZZ1 chemical tag could enhance electrostatic interaction with the YPEL5 basic groove (Fig. 5a). To test this idea, we synthesized ZZ2, which has an additional chloro substituent adjacent to the existing sulfonyl fluoride moiety of ZZ1 (Fig. 5b). In doing so, we engaged in rational structure-based improvement of a fortuitously discovered c-Glue.

**Figure 5:**
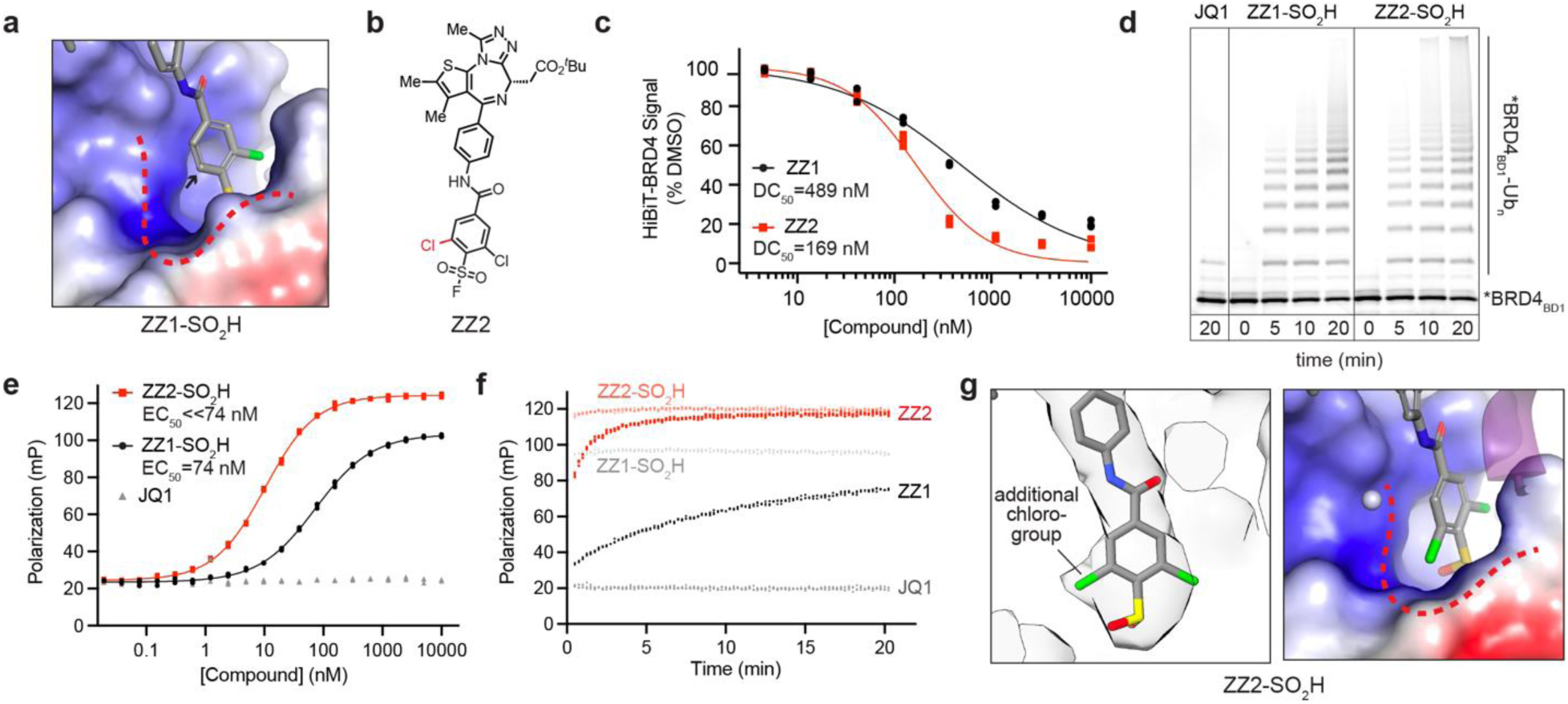
Structure-based improvement of the c-Glue. a) Electrostatic potential surface of the ZZ1-SO_2_H-bound YPEL5 groove showcasing a vacant basic pocket (outlined with a red dash) adjacent to the unsubstituted position of the degrader’s chemical tag (indicated by an arrow). b) Chemical structure of ZZ2 highlighting the incorporated chloro group (red) at the second *ortho*-position of ZZ1’s chemical tag. c) HiBiT-BRD4 assay results for Jurkat cells treated with the indicated compounds for 5 h. d) Qualitative comparison of ZZ1-SO_2_H and ZZ2-SO_2_H ability to trigger *in vitro* YPEL5-GID/CTLH E3-catalyzed *BRD4_BD1_ ubiquitylation. ZZ2-SO_2_H was generated by pre-incubation of ZZ2 in the DTT-containing buffer (Extended Data Figure 6d). e) FP assay quantifying propensity of ZZ1-SO_2_H and ZZ2-SO_2_H to promote ternary complex formation. Note that polarization values obtained upon ZZ2-SO_2_H titration fit the “[agonist] vs. response” model but its superior MGD activity precludes accurate estimation of EC_50_. f) Real-time FP assay testing rates of ternary complex formation triggered by ZZ1 and ZZ2 as well as their acidic derivatives. Fluorescence polarization was monitored over time upon mixing *BRD4_BD1_ and YPEL5-WDR26 with different versions of the degraders. g) Close-up of the ZZ2-SO_2_H-induced ternary complex cryo-EM structure, resolved to 3.4 Å and sharpened with DeepEMhancer. The images highlight the overall fit of the degrader’s chemical tag (left) and the filling of the vacant YPEL5 basic pocket by the introduced chloro group according to the structure-based design (right).

ZZ2 is a superior BRD4 degrader when compared with ZZ1 (ZZ2 DC_50_ = 169 nM; ∼3-fold more potent than ZZ1), and it retains degradation specificity for BET proteins (Fig. 5c, Extended Data Fig. 6c). In line with this improved degradation potency, ZZ2-SO_2_H promotes more robust BRD4_BD1_ ubiquitylation and mediates a more stable ternary complex manifested by higher fluorescence polarization at saturating degrader concentration and substantially lower EC_50_ (Fig. 5d, 5e, Extended Data Fig. 6d). Moreover, ZZ2 induces the BRD4-compound-YPEL5 ternary complex at a significantly faster rate *in vitro* when compared with ZZ1 (Fig. 5f). We surmise the rate enhancement is the consequence of two factors: improved YPEL5 engagement and accelerated conversion of the sulfonyl fluoride moiety to the active sulfinic acid, facilitated by the electron-withdrawing chloro group. In agreement with the structure-based design, the 3.4-Å cryo-EM structure of the ZZ2-SO_2_H-mediated ternary complex showed an extra portion of electron density corresponding to the new chloro substituent occupying the targeted YPEL5 pocket (Fig. 5g, Extended Data Table 1, Extended Data Fig. 7). Aside from the added chloro substituent and its contacts, ZZ2-SO_2_H largely maintains the interactions observed in the ZZ1-SO_2_H-induced complex (aligning with RMSD=0.3 Å) (Extended Data Fig. 6e). Therefore, ZZ2 substantiates the importance of electrostatic anchoring of the negatively charged c-Glue in the basic groove of YPEL5.

## Discussion

**ZZ1** is a GID/CTLH-dependent BRD4 MGD that uses a prodrug mechanism to expose a negatively charged ligase-binding moiety upon its entry into target cells. Cryo-EM structures of the neosubstrate recognition complex show that, once exposed, a negatively charged sulfinic acid moiety interacts with a basic pocket on YPEL5, a GID/CTLH subunit essential to most cancer cell lines^42^. Sulfinic acid perception is a central feature of the recognition complex, and we therefore designate **ZZ1** a c-Glue. The structures also demonstrate structural homology between YPEL5 and the thalidomide-binding domain of the IMiD (Immunomodulatory Drug) MGD target ubiquitin ligase, CRBN. Contrary to its only known biochemical ability, which is to inhibit a GID/CTLH-substrate interaction^43^, YPEL5 acts as a substrate receptor in **ZZ1**-treated cells. Rational design and synthesis of the improved compound, **ZZ2**, demonstrates that these properties can be exploited to create more potent YPEL5-dependent c-Glues by structure-guided medicinal chemistry.

**ZZ1** and **ZZ2** are c-Glues that represent a new strategy for charge-driven proximity pharmacology. Although these compounds are MGDs like the well-known IMiDs, three features set them apart. First, **ZZ1** and **ZZ2** undergo intracellular chemical activation and, upon conversion, become MGDs that induce a noncovalent and cooperative BRD4-compound-YPEL5 ternary complex. Second, the new compounds use charged interactions rather than more traditional hydrophobic small molecule-protein contacts. Indeed, the CRBN-IMiD binding site is a hydrophobic “tri-tryptophan pocket.” Third, neither new compound appreciably binds YPEL5 in the absence of their bromodomain-containing targets. This final characteristic is a key distinction; IMiD drugs bind an E3 ligase (CRBN) and cannot bind neosubstrates in its absence. The switch from E3 ligase to target as the primary binding partner indicates that future MGD discovery campaigns may be initiated “target-side,” enabling greater control over eventual MGD pharmacology. Recent discovery of covalent MGDs supports this idea, and the work we report here formalizes this strategy and eliminates the need for covalency^30,48-50^. The c-Glues we describe herein are *bona fide* MGDs and retain their potency even at high compound concentrations, bypassing the so-called “hook effect,” which is a common limitation of PROTACs and covalent degraders^30^. These findings demonstrate that the GID/CTLH E3 ubiquitin ligase complex is amenable to charge-driven induced proximity, similar to established clinical small molecule MGDs targeting CRLs.

The discovery of YPEL5 as a MGD receptor was unexpected, since its only documented function is to inhibit ubiquitylation of NMNAT1 (metabolic enzyme nicotinamide/nicotinic-acid-mononucleotide-adenylyltransferase 1)^43^. YPEL5 and NMNAT1 compete for an overlapping binding site on another GID/CTLH E3 subunit, WDR26. Whether YPEL5 is an endogenous E3 ligase substrate receptor will require identification of its native substrates, and the cryo-EM structures presented here strongly suggest these will feature acidic degrons. Nevertheless, our detailed study of YPEL5 demonstrates unexpected functional plasticity for ubiquitin ligases; an inhibitor of substrate ubiquitylation for one set of targets can stimulate the ubiquitylation of a second set. The versatility of the GID/CTLH E3 system showcases this concept^39,40,43,51^.

GID/CTLH complexes are modular, and multiple auxiliary subunits are reported substrate receptors. These include GID4^39,52,53^, WDR26^40,43^, FAM72A^54^, and, as now demonstrated in the context of a MGD, YPEL5. Diverse mechanisms of substrate recognition presumably facilitate the recruitment and positioning of a wide range of native substrates and, consequently, neosubstrates when presented with dedicated MGD compounds. Furthermore, the two-fold symmetric architecture of GID/CTLH (Fig. 2a) exposes asymmetric neosubstrates to two distinct ubiquitylation active sites poised for ubiquitin transfer to distinct sets of neosubstrate lysine side chains. Future YPEL5-dependent small molecule degraders could be developed to enable a multimeric neosubstrate to engage two substrate-binding modules with enhanced avidity^39^. Avid neosubstrate binding would enable meaningful TPD pharmacology even for relatively weak ternary complex interactions.

Prodrugs offer a compelling strategy for targeted disease therapy^55,56^. Chemotherapeutics like capecitabine and irinotecan are now classic examples of tumor cell-specific metabolic prodrug activation^57^, but this strategy is not common in TPD aside from a handful of covalent degraders^48-50^ and engineered PROTACs^58-62^. Sulfonyl fluoride chemistry, which is not a conventional feature of prodrugs, confers favorable c-Glue characteristics when activated by electron-deficient groups.

The first characteristic is cell permeability of the prodrug but not the activated compound. The second characteristic is dependence of prodrug conversion on reducing equivalents. This prevents premature extracellular activation and may confer selective activity in tumor cells, which harbor elevated GSH levels^63^. Using cancer-specific reactive metabolites to activate prodrugs presents a compelling strategy that may enable targeted accumulation of MGDs in disease tissues and could further expand the therapeutic index enabled by tissue-specific ubiquitin ligase expression. The third and most important characteristic, definitional to c-Glue, is the ability to target a positively charged protein cavity. In principle, the same c-Glue strategy can be used to target related E3 ligase families with positively charged pockets that accommodate endogenous phosphodegrons (*e.g.* F-box and SOCS-box proteins)^20,21^ or protein C-terminal acids (*i.e.* C-degron pathway E3 ligases)^22-24^.

In addition to the possibility of applying the c-Glue concept to new small molecule degraders, the structural similarity between YPEL5 and CRBN suggests direct access to a MGD discovery strategy analogous to what has been done for the IMiDs. Chemical diversification of a CRBN-binding core has produced a plethora of TPD therapeutics that are now entering clinical trials^2,14^. Similarly, improved YPEL5 ligands from the structure-based medicinal chemistry campaign we have initiated will enable chemocentric MGD discovery via elaboration of a YPEL5-binding chemical core. Such libraries would be intrinsically limited in their activities to cells with pre-existing high YPEL5 protein levels – YPEL5 is overexpressed in developing erythrocytes and Ewing Sarcomas^42,43^ – providing a compelling path to a broad therapeutic window. Indeed, the GID/CTLH complex is an attractive candidate TPD effector due to its frequent upregulation in cancer and broad essentiality^64,65^. The latter characteristic will presumably prevent the rapid emergence of ligase loss-driven resistance. CRBN mimicry by the YPEL5-GID/CTLH ubiquitin ligase, enabled by the development of c-Glues, provides a blueprint for next-generation TPD therapeutics discovery.

**Extended Data Figure 1:**
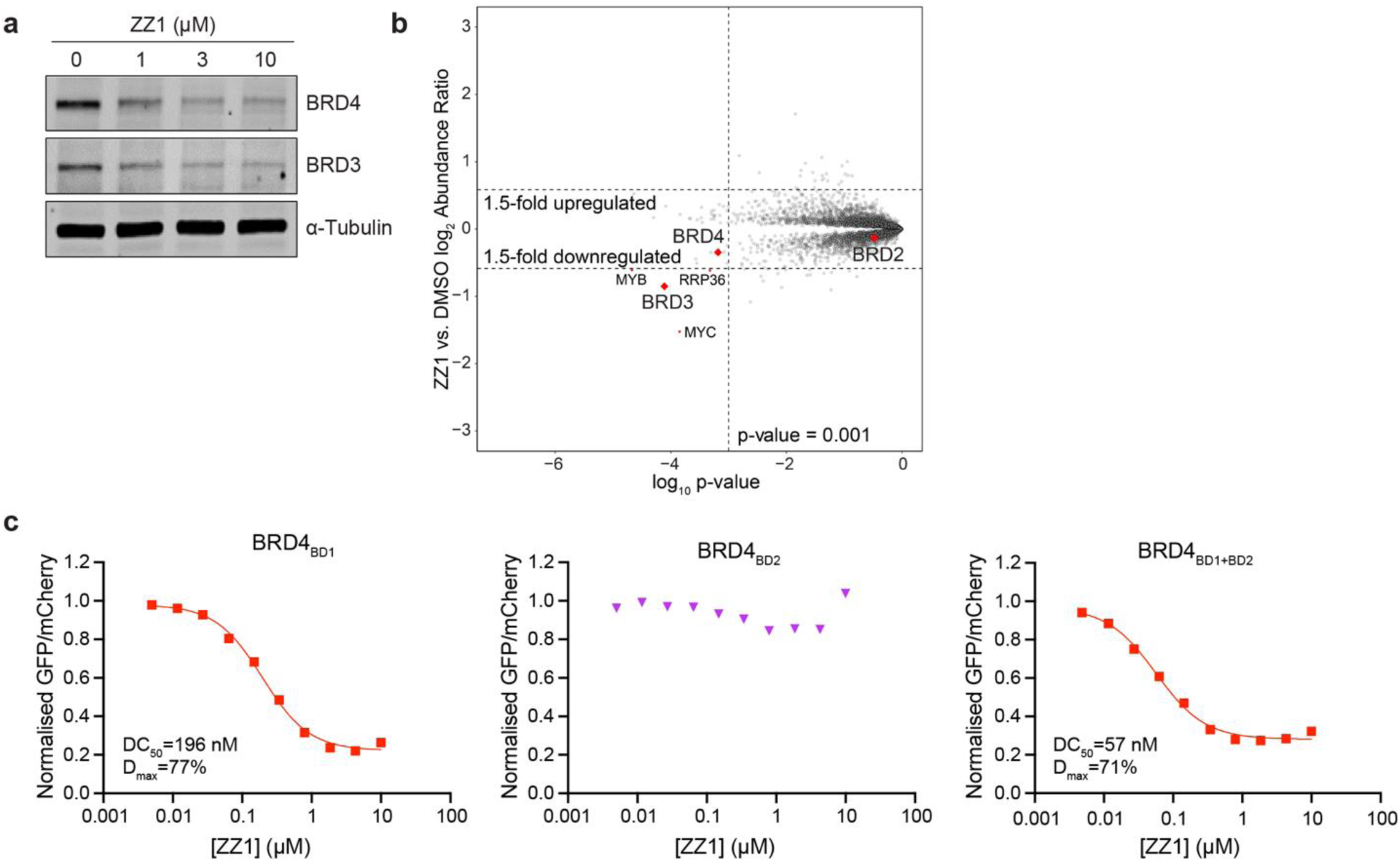
Determining target selectivity of ZZ1-induced degradation. a) Western blots showing BRD4 and BRD3 degradation in MOLT-4 cells after 5 h treatment with ZZ1. b) Quantitative proteome-wide mass spectrometry in MOLT-4 cells after 3 h treatment with 1 µM of ZZ1. c) Identifying of the BRD4 region required for ZZ1-induced degradation with a cellular fluorescent reporter assay. The examined reporters were either the isolated BRD4 bromodomains (BD1 or BD2) or a tandem construct comprising BD1 and BD2 connected by the intervening native sequence (BRD4_BD1+BD2_).

**Extended Data Figure 2:**
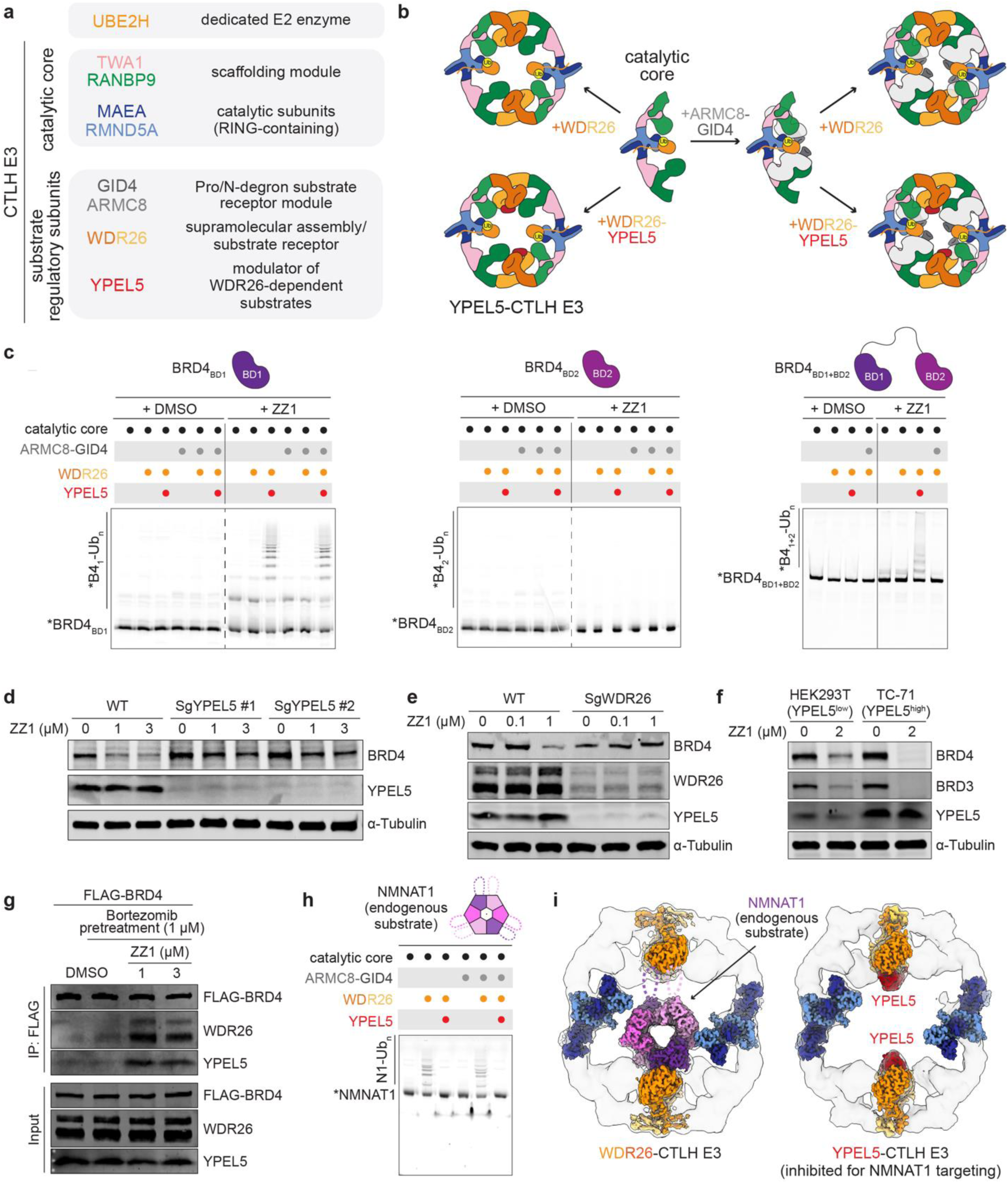
YPEL5-GID/CTLH E3 ligase is an effector for ZZ1-mediated BRD4 targeting. a) Color-coded guide to the GID/CTLH E3 subunits and their reported functions. b) Schematic illustrating the architecture of the GID/CTLH E3 ligases, which share a common catalytic core that associates with divergent auxiliary subunits enabling substrate targeting. WDR26 acts as a supramolecular assembly factor by connecting two copies of the catalytic core (either alone or bound to GID4-ARMC8) into a singular giant oval structure with a large hollow center. Each WDR26 homodimer in the supramolecular assembly can bind a single copy of YPEL5, yielding the YPEL5-GID/CTLH E3. Subunits are colored according to the guide in (a). c) Identifying E3 ligase leveraged by ZZ1 with *in vitro* ubiquitylation assays. The suite of GID/CTLH E3 assemblies shown in (b) was tested for activity towards fluorescent BRD4 bromodomain substrates, either in isolation (BRD4_BD1_ and BRD4_BD2_) or in tandem (BRD4_BD1+BD2_). Asterisk denotes the fluorescent FAM label appended to substrates’ N-termini. All reactions were quenched after 45 min. d) Western blots showing BRD4 degradation in WT or YPEL5-KO Jurkat cells treated with the indicated concentration of ZZ1 for 5 h. e) Western blots showing BRD4 degradation in WT or WDR26-KO HEK293T cells treated with the indicated concentration of ZZ1 for 5 h. f) Western blots showing BRD4 degradation in HEK293T (YPEL5*^low^*) or TC-71 (YPEL5*^high^*) cells treated with the 2 µM of ZZ1 for 5 h. g) Co-immunoprecipitation of FLAG-tagged BRD4 and YPEL5-WDR26-containing GID/CTLH E3 in the presence of ZZ1. FLAG-tagged BRD4 transfected cells were preincubated with the proteasomal pathway inhibitor (bortezomib) for 1 h to prevent BRD4 degradation. h) *In vitro* ubiquitylation assay as in (b) but performed with the endogenous NMNAT1 substrate to recapitulate its previously reported GID/CTLH E3-dependent regulation. In contrast to its essential role in ZZ1-induced BRD4 ubiquitylation, YPEL5 acts as an inhibitor of NMNAT1 targeting. Reactions were quenched after 45 min. i) Previous cryo-EM structures of NMNAT1- and YPEL5-bound GID/CTLH E3 assemblies (EMD-18175 and EMD-18170, respectively)^43^ fit with segmented focused-refined maps of relevant modules (EMD-18345 and EMD-18316, respectively) explaining the biochemically defined mode of NMNAT1 targeting: the hexameric NMNAT1 and YPEL5 are engaged by the overlapping binding sites on dimeric WDR26 modules. Consequently, YPEL5 sterically blocks WDR26-mediated NMNAT1 recruitment.

**Extended Data Figure 3:**
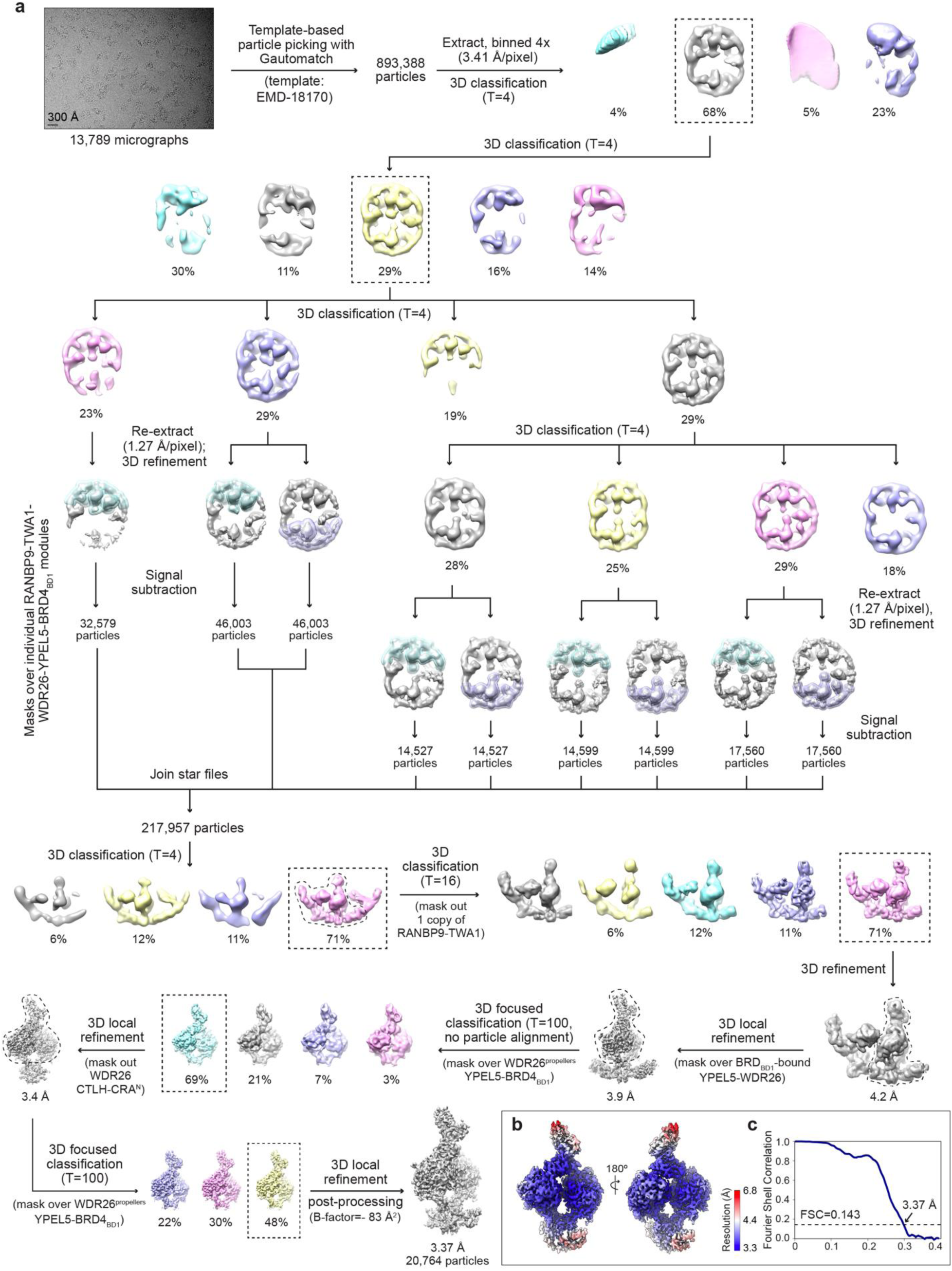
Determination of the cryo-EM structure of the ZZ1-SO_2_H-induced ternary complex. a) Flowchart of the cryo-EM data processing workflow generating the focused-refined map of the ternary complex comprising ZZ1-SO_2_H c-Glue, YPEL5-WDR26 E3 receptor module and BRD4_BD1_ neosubstrate. The scale bar in the motion-corrected representative micrograph corresponds to 300 Å. b) The final post-processed map color-coded to illustrate variations in its local resolution. c) Gold-standard Fourier shell correlation (FSC) plot. The dotted line represents 0.143 cut-off criterion for estimating nominal resolution.

**Extended Data Figure 4:**
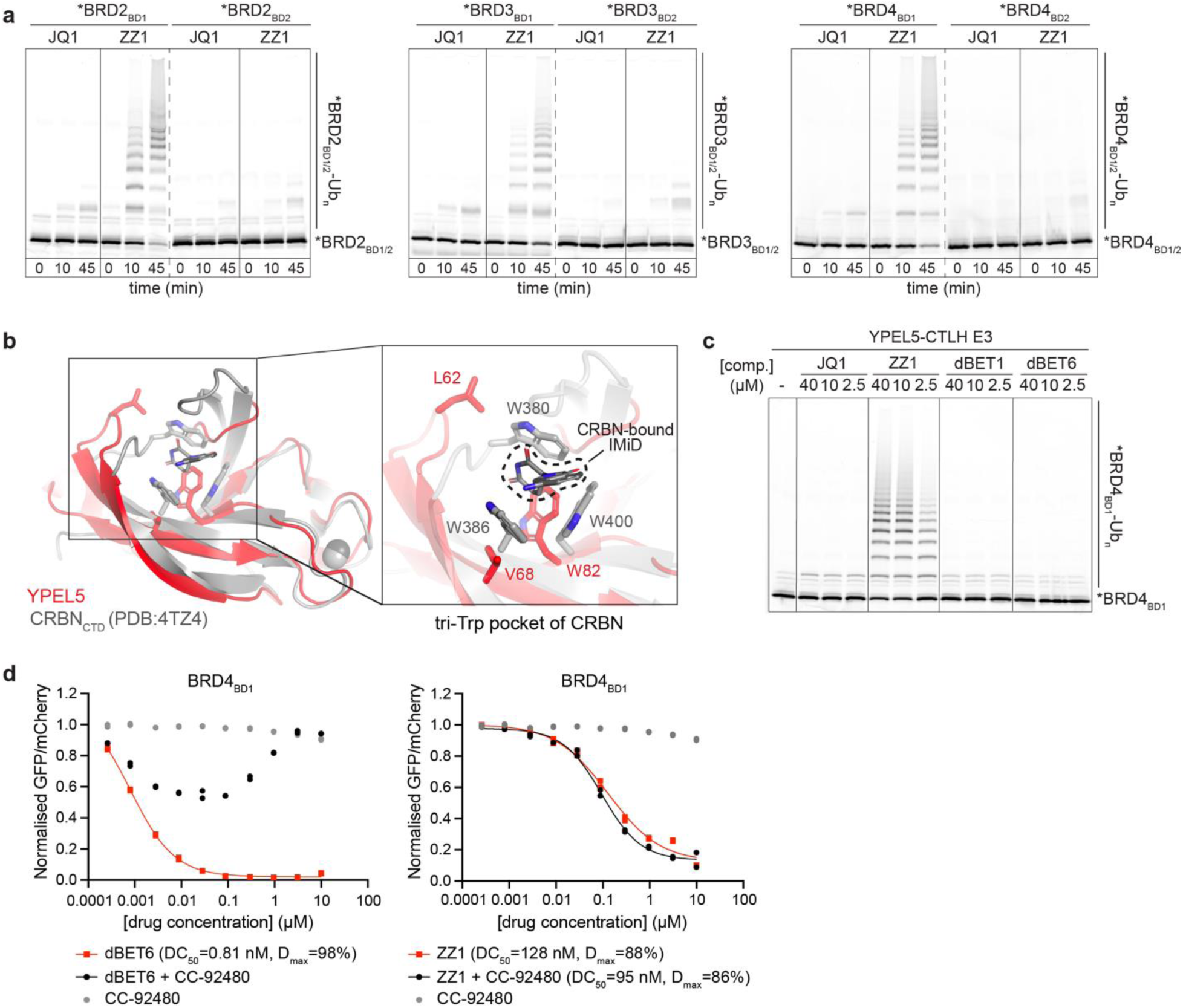
Identifying neosubstrate and degrader specificity of YPEL5. a) *In vitro* ubiquitylation assays testing specificity of ZZ1-induced ubiquitylation towards bromodomains of BET protein family members that all bind the parental substrate-recruiting JQ1 handle of ZZ1. b) Superposition of the YPEL5 structure with that of the thalidomide binding domain of CRBN (CRBN_CTD_) engaging an Immunomodulatory Drug (IMiD) MGD (PDB: 4TZ4). The close-up highlights CRBN residues forming the IMiD-engaging hydrophobic site (the “tri-Trp pocket”) and the corresponding YPEL5 residues (shown as sticks). Despite adopting a homologous fold, YPEL5 does not possess two out of three residues critical for IMiD binding. c) Examining ligand-binding preference of YPEL5 by performing *in vitro* BRD4_BD1_ ubiquitylation assay with ZZ1 and CRBN-based PROTACs employing the BRD4 ligand JQ1. d) Analysis of BRD4_BD1_-eGFP degradation in K562 stability reporter cells treated with indicated compounds or compound combinations (co-treatment with equimolar mixtures).

**Extended Data Figure 5:**
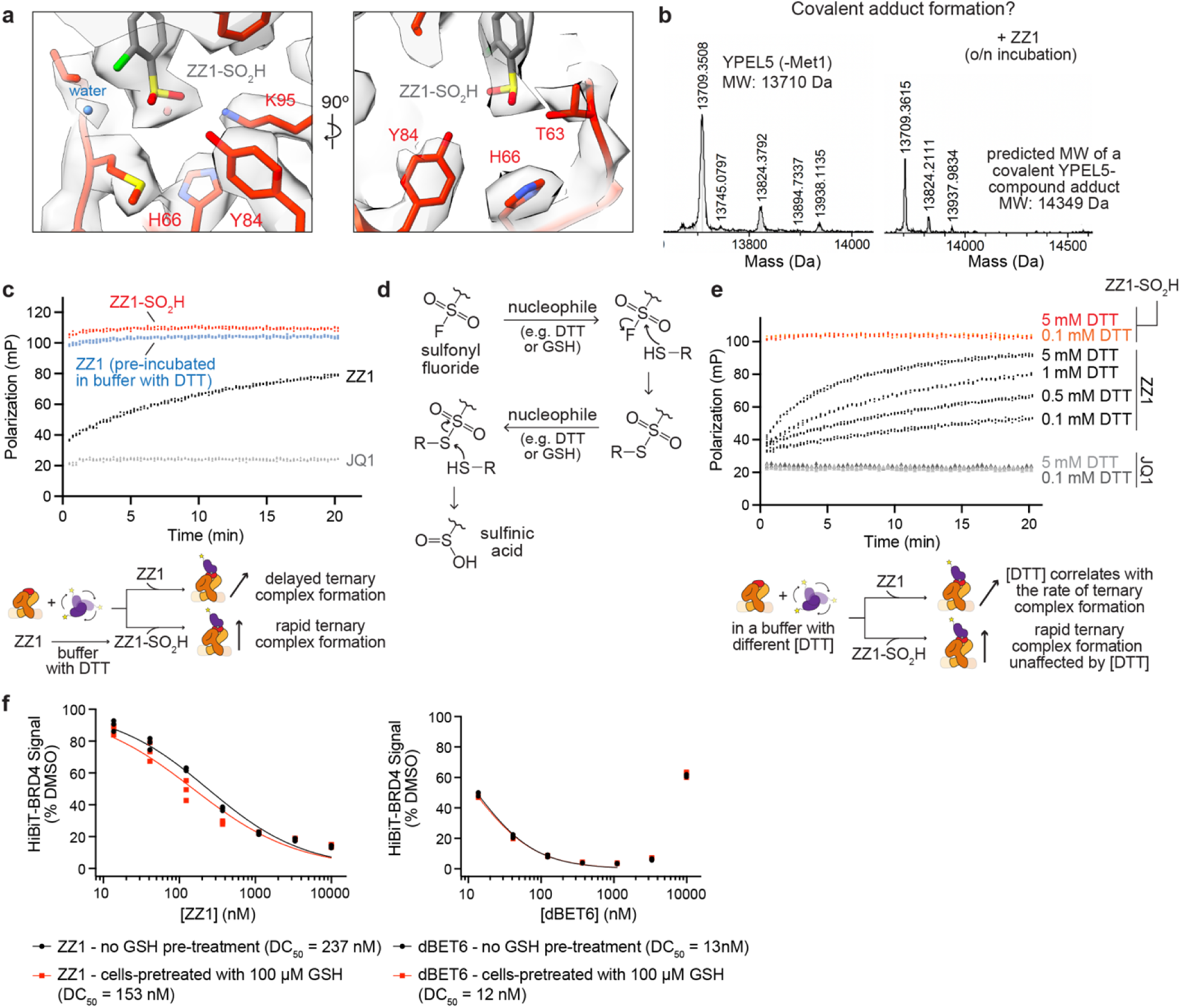
Biochemical and structural assays probing ZZ1 mode-of-action. a) Close-up of electron density (gray transparent) in the cryo-EM structure of the ternary complex (shown in Figure 2b) corresponding to the ZZ1-SO_2_H chemical tag and the surrounding YPEL5 residues, along with their atomic coordinates (sticks). Absence of continuous density between the sulfinic acid and neither YPEL5 nucleophilic amino acid side chains suggests the non-covalent ZZ1 mode-of-action. b) Intact mass spectrometry analysis testing formation of potential ZZ1-induced covalent adducts between YPEL5 (within the YPEL5-WDR26 subcomplex) and BRD4_BD1_. c) Real-time FP assay probing kinetics of ternary complex formation in the DTT-containing buffer induced by ZZ1, ZZ1-SO_2_H, or ZZ1 pre-incubated in the FP buffer. Polarization signal was measured over time after combining the protein mix (*BRD4_BD1_ and YPEL5-WDR26) prepared in the DTT-containing buffer with the degrader compounds. d) Schematic of the proposed mechanism of sulfonyl fluoride conversion to sulfinic acid triggered by nucleophilic thiol groups of reducing agents, such as DTT (*in vitro*) or GSH (in cells). e) Real-time FP assay testing effect of DTT on the rate of the ZZ1-SO_2_H-induced ternary complex formation. The varying DTT concentrations during the experiment were controlled by the composition of the buffer used for preparing the protein mix (*BRD4_BD1_ and YPEL5-WDR26) prior to degrader addition. f) HiBiT-BRD4 assay results for Jurkat cells pre-treated with 100 µM GSH for 2 h, followed by treatment with the indicated compounds for 5 h.

**Extended Data Figure 6:**
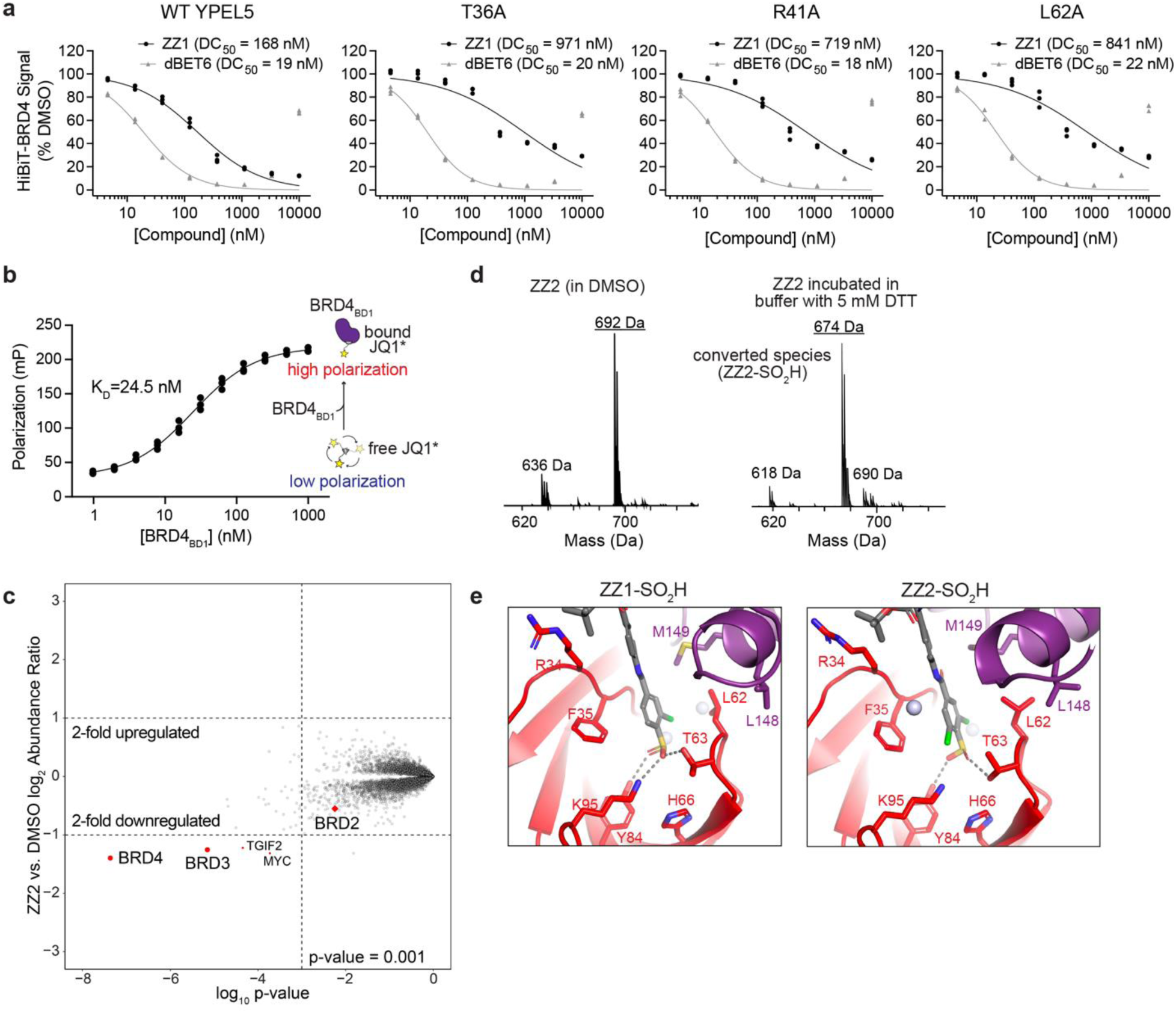
Mechanistic insights into ZZ1 and ZZ2 c-Glue activity. a) HiBiT-BRD4 assay results for WT and mutant YPEL5-expressing Jurkat cells treated with the indicated compounds for 5 h. The plots comparing the estimated DC_50_ values are presented in Figure 4d. b) FP assay for establishing the competitive ligand displacement experiment probing cooperative ternary complex formation (Figure 4e). Polarization values upon BRD4_BD1_ titration to the fluorescent JQ1 tracer (FAM-JQ1) were fit to the one-site binding model to estimate the BRD4_BD1_-JQ1 affinity (equilibrium dissociation constant, K_D_). c) Quantitative proteome-wide mass spectrometry in MOLT-4 cells after 3 h treatment with 1 µM ZZ2. d) Intact mass spectrometry demonstrates conversion of the sulfonyl fluoride moiety of ZZ2 (left) to sulfinic acid (right) after incubation in a DTT-containing buffer. Peaks corresponding to lower molecular weight species correspond to ZZ2 and ZZ2-SO_2_H derivatives formed upon hydrolysis of their *tert*-butyl ester. e) Close-up of ternary complex structures induced by ZZ1-SO_2_H and ZZ2-SO_2_H highlighting their common binding mode. YPEL5 and BRD4 residues involved in protein-protein and/or protein-degrader contacts are shown as sticks. Spheres represent water molecules, while dashes denote hydrogen bonds.

**Extended Data Figure 7:**
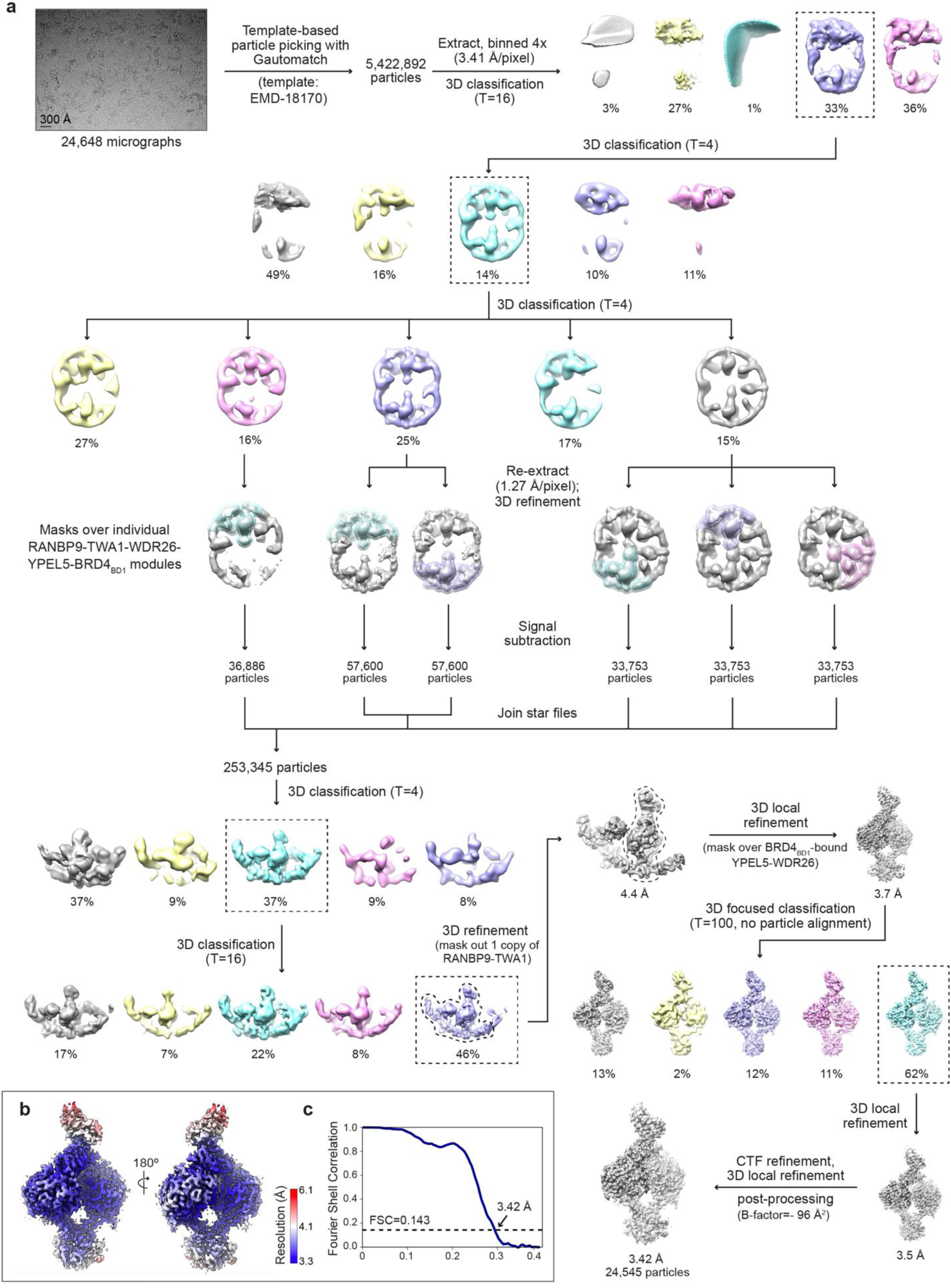
Determination of the cryo-EM structure of the ZZ2-SO_2_H-induced ternary complex. a) Flowchart of the cryo-EM data processing workflow generating the focused-refined map of the ternary complex comprising the improved ZZ2-SO_2_H c-Glue, YPEL5-WDR26 E3 receptor module and BRD4_BD1_ neosubstrate. The scale bar in the motion-corrected representative micrograph corresponds to 300 Å. b) The final post-processed map color-coded to illustrate variations in its local resolution. c) Gold-standard Fourier shell correlation (FSC) plot. The dotted line represents 0.143 cut-off criterion for estimating nominal resolution.

**Extended Data Table 1.**
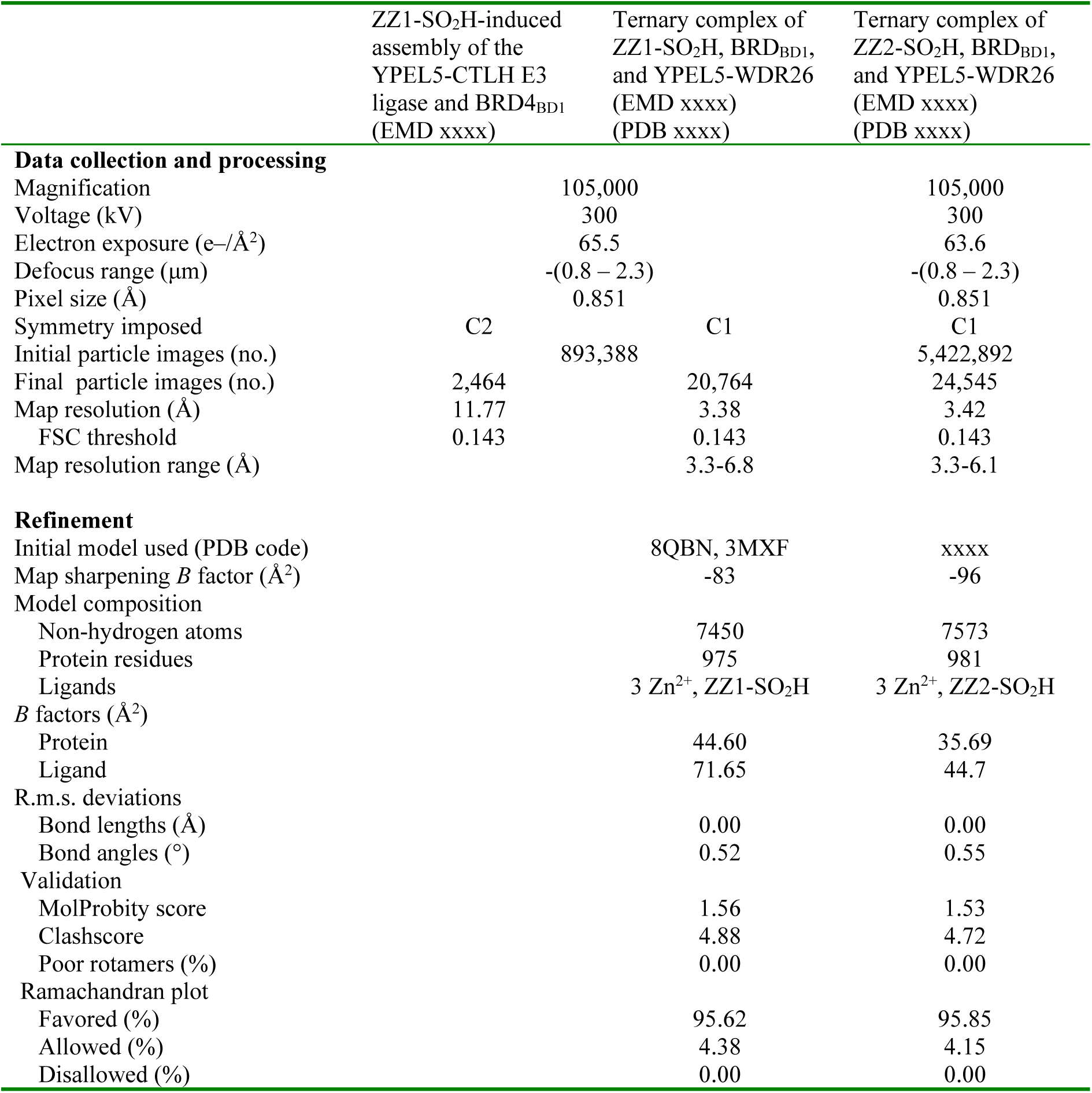
Cryo-EM data collection, refinement and validation statistics.

## Methods

### Chemical synthesis

Additional details are provided in the supporting information.

### General cell biology methods

Roswell Park Memorial Institute (RPMI) 1640 medium and Dulbecco’s modified Eagle’s medium (DMEM), Iscove’s Modified Dulbecco’s Medium (IMDM), Fetal bovine serum (FBS), penicillin– streptomycin (10,000 units/mL sodium penicillin G and 10,000 μg/mL streptomycin), trypsin– EDTA solution (1×), and phosphate-buffered saline (PBS; 1×) were purchased from Gibco Invitrogen Corp. (Grand Island, NY, USA). MG132, MLN-4924, Bortezomib, and dBET6 were purchased from MedChemExpress (Monmouth Junction, NJ, USA). All other chemicals were purchased from Sigma-Aldrich (St. Louis, MO, USA), unless indicated otherwise.

### Mammalian cell culture

Human leukemia (Jurkat and MOLT-4) and a kidney epithelial (HEK293T) cell lines were obtained from the American Type Culture Collection (ATCC, Manassas, VA, USA). The K562-Cas9 was provided by Z. Tothova (Dana-Farber Cancer Institute), and the Ewing sarcoma cell line (TC-71) was obtained from R. George (Dana-Farber Cancer Institute). Cells were cultured in medium (RPMI 1640 for Jurkat, MOLT-4, K562 cells; DMEM medium for HEK293T cells; IMDM for TC-71 cells) supplemented with 10% heat-inactivated FBS, 100 units/mL penicillin, 100 µg/mL streptomycin, and 0.25 µg/mL amphotericin B. Cells were incubated at 37 °C with 5% CO_2_ in a humidified atmosphere. Mycoplasma testing was performed monthly using the MycoAlert mycoplasma detection kit (Lonza, Basel, Switzerland) and all lines were negative.

### Generation of HiBiT-BRD4 cells

Introduction of a HiBiT coding sequence into the endogenous BRD4 locus in Jurkat cells was achieved using CRISPR-Cas9 genome editing. Alt-R^TM^ CRISPR RNA (crRNA) and trans-activating CRISPR RNA (tracrRNA) (Integrated DNA Technologies, Coralville, IA, USA) were resuspended in Nuclease-Free Duplex Buffer (Integrated DNA Technologies) at a concentration of 200 µM each. Equal volumes of crRNA and tracrRNA were mixed (final concentration of 100 µM each) and heated for 5 minutes at 95 °C. After heating, the complex was gradually cooled to room temperature. The oligo complex was then incubated at room temperature for 20 minutes with the Alt-R^TM^ Cas9 Nuclease V3 (Integrated DNA Technologies) to form the ribonucleoprotein (RNP) complex. The double-stranded DNA HDR template (HiBiT-BRD4 with extensions, see below), the RNP complex, and an electroporation enhancer (Integrated DNA Technologies) were then electroporated into Jurkat cells using an Amaxa^TM^ 4D-Nucleofector (Lonza). Electroporated cells were transferred to medium with HRD enhancer (Integrated DNA Technologies). Single cells were subsequently isolated via fluorescence activated cell sorting, and HiBiT expression from individual clones was detected with the Nano-Glo HiBiT Lytic Detection System (Promega, Madison, WI, USA). Sequences for crRNA and HDR doner are provided below.

>crRNA sequence

ACTAGCATGTCTGCGGAGAG

>HDR donor sequence (+)

CATTACTGGCAGATTTCTCAATCTCGTCCCAGGGCCGCTCTCCGCAGAGCCAGAACTCCCGCTAATCTTCTTGAACAGCCGCCAGCCGCTCACCATGCTAGTGATCCCATCACATTCTTCACCAGGCACTCTA

>HDR donor sequence (-)

TAGAGTGCCTGGTGAAGAATGTGATGGGATCACTAGCATGGTGAGCGGCTGGCGGCTGTTCAAGAAGATTAGCGGGAGTTCTGGCTCTGCGGAGAGCGGCCCTGGGACGAGATTGAGAAATCTGCCAGTAATG

### HiBiT-BRD4 assay

Endogenous BRD4 protein levels were evaluated using the Nano-Glo HiBiT Lytic Detection System (Promega). In brief, 1.5 × 10^4^ HiBiT-BRD4 Jurkat cells were seeded into 384-well plates and incubated with the indicated concentrations of compounds. After 5 h, the plates were subjected to Nano-Glo HiBiT Lytic Detection System as described in manufacturer’s manual. The HiBiT-BRD4 assays were conducted in biological triplicates. IC_50_ values were determined using a non-linear regression curve fit in GraphPad PRISM v10.3.1.

### Reporter vectors

The Cilantro 2 reporter vector (PGK.BsmBICloneSite-10aaFlexibleLinker-eGFP.IRES.mCherry. cppt.EF1α.PuroR, Addgene #74450) was used for flow-based degradation assays. The following boundaries for BRD4 truncations were used: BRD4_BD1_: (UniProt entry O60885, residues 14-138) or BRD4_BD2_: (residues 306-418). Reporter cell lines were generated as described before.^30^

### BRD4 reporter stability analysis

K562-Cas9 cells expressing the BRD4_BD1_-eGFP, BRD4_BD2_-eGFP, or BRD4_BD1+BD2_-eGFP degradation reporters were resuspended at 0.7 × 10^6^ ml^−1^. 50 µl of cell suspension was seeded in 384-well plates and immediately treated with DMSO or drug for 16 h. The indicated drugs were dispensed with a D300 digital dispenser (Tecan Genomics, Männedorf, Switzerland). The fluorescent signal was quantified by flow cytometry (FACSymphony flow cytometer, BD Biosciences, Franklin Lakes, NJ, USA). Using FlowJo (Flow cytometry analysis software, BD Biosciences), the geometric mean of the eGFP and mCherry fluorescent signal for round and mCherry-positive cells was calculated. The ratio of eGFP to mCherry was normalized to the average of ten DMSO-treated controls.

### Bison CRISPR screen for BRD4 stability

The Bison CRISPR library targets 713 E1, E2 and E3 ubiquitin ligases, deubiquitinases and control genes and contains 2,852 guide RNAs (Addgene #169942)^69^. 10% (v/v) of the Bison CRISPR library was added to 10 × 10^6^ BRD4(BD1)_eGFP_ or BRD4(BD2)_eGFP_ K562-Cas9 cells and transduced (2,400 rpm, 2 h, 37 °C). Eight days later, cells were treated with drug or DMSO for 16 h and four populations were collected (top 5%, top 5–15%, lowest 5–15% and lowest 5%) on the basis of the BRD4_eGFP_ to mCherry mean fluorescent intensity (MFI) ratio on an MA900 Cell Sorter (Sony, Minato City, Japan). Sorted cells were collected by centrifugation, subjected to direct lysis, and amplified as described before.^28^ Amplified sgRNAs were quantified using the Illumina NovaSeq SP platform (Genomics Platform, Broad Institute). The screen was analyzed by comparing stable populations (top 5% eGFP/mCherry expression) to unstable populations (lowest 5% eGFP/mCherry expression) as described before.^28^ Note that, except for RMND5B, the statistically significant hits were the only genes coding for GID/CTLH-related proteins targeted by the Bison library. We speculate that the insensitivity of BRD4 degradation to RMND5B KO is due to its paralog, RMND5A (not targeted by the library) playing a redundant role or the two proteins being involved in different suites of GID/CTLH E3 complexes, with those containing RMND5A being responsible for ZZ1-dependent activity.

### Western blotting analysis

Total cells lysates were prepared in 2× sample loading buffer (i.e., 250 mM Tris-hydrochloride: pH 6.8, 4% sodium dodecyl sulfate, 10% glycerol, 0.006% bromophenol blue, 2% β-mercaptoethanol, 50 mM sodium fluoride, and 5 mM sodium orthovanadate). The samples with cell lysates were boiled for 5-8 min at 95 °C. The protein concentrations of the cell lysates were quantified using the BCA method and a BCA Protein Assay Kit (Thermo Fisher Scientific, Waltham, MA, USA). Equal amounts of protein were subjected to 4-20% sodium dodecyl sulfate-polyacrylamide gel electrophoresis and transferred to nitrocellulose membranes (cat. no. 1620112; Bio-Rad Laboratories, Hercules, CA, USA). The membranes were blocked using Intercept^®^ (TBS) Blocking Buffer (LI-COR Biosciences, Lincoln, NE, USA), and subsequently probed with appropriate primary antibodies [anti-BRD4 (cat. no. ab128874; Abcam, Cambridge, UK and cat. no. 63759; Cell Signaling Technology, Danvers, MA, USA), anti-α-Tubulin (cat. no. 3873; Cell Signaling Technology), anti-YPEL5 (cat. no. PA5-26957; Invitrogen), anti-WDR26 (cat. no. NBP1-83628; Novus Biologicals, Centennial, CO, USA), anti-BRD3 (cat. no. ab50818, Abcam), anti-FLAG M2 (cat. no. F1804; Sigma-Aldrich)] at 4 °C overnight and then incubated with IRDye 800-labeled goat anti-rabbit IgG (LI-COR Biosciences, cat. no. 926-32211) or IRDye 680RD goat anti-Mouse IgG (LI-COR Biosciences, cat. no. 926-68070) secondary antibodies at room temperature for 1 h. After washing the membranes with PBS for 30 min, the membranes were detected on Li-COR Odyssey CLx system.

### Sample preparation for quantitative LFQ quantitative mass spectrometry

Cells were lysed by addition of lysis buffer (8 M Urea, 50 mM NaCl, 50 mM 4-(2-hydroxyethyl)-1-piperazineethanesulfonic acid (EPPS) pH 8.5, Protease and Phosphatase inhibitors) and homogenization by bead beating (BioSpec, Bartlesville, OK, USA) for three repeats of 30 seconds at 2400 strokes/min. Bradford assay was used to determine the final protein concentration in the clarified cell lysate. Fifty micrograms of protein for each sample was reduced, alkylated and precipitated using methanol/chloroform as previously described^70^ and the resulting washed precipitated protein was allowed to air dry. Precipitated protein was resuspended in 4 M urea, 50 mM HEPES pH 7.4, followed by dilution to 1 M urea with the addition of 200 mM EPPS, pH 8. Proteins were digested with the addition of LysC (1:50; enzyme:protein) and trypsin (1:50; enzyme:protein) for 12 h at 37 °C. Sample digests were acidified with formic acid to a pH of 2-3 before desalting using C18 solid phase extraction plates (SOLA, Thermo Fisher Scientific). Desalted peptides were dried in a vacuum-centrifuged and reconstituted in 0.1% formic acid for liquid chromatography-mass spectrometry analysis.

Data were collected using a TimsTOF HT (Bruker Daltonics, Bremen, Germany) coupled to a nanoElute LC pump (Bruker Daltonics) via a CaptiveSpray nano-electrospray source. Peptides were separated on a reversed-phase C_18_ column (25 cm x 75 µm ID, 1.6 µM, IonOpticks, Fitzroy, Australia) containing an integrated captive spray emitter. Peptides were separated using a 50 min gradient of 2 - 30% buffer B (acetonitrile in 0.1% formic acid) with a flow rate of 250 nL/min and column temperature maintained at 50 °C.

The TIMS elution voltages were calibrated linearly with three points (Agilent ESI-L Tuning Mix Ions; 622, 922, 1,222 *m/z*) to determine the reduced ion mobility coefficients (1/K_0_). To perform diaPASEF, we used py_diAID^71^, a python package, to assess the precursor distribution in the *m/z*-ion mobility plane to generate a diaPASEF acquisition scheme with variable window isolation widths that are aligned to the precursor density in m/z. Data was acquired using twenty cycles with three mobility window scans each (creating 60 windows) covering the diagonal scan line for doubly and triply charged precursors, with singly charged precursors able to be excluded by their position in the m/z-ion mobility plane. These precursor isolation windows were defined between 350 - 1250 *m/z* and 1/k0 of 0.6 - 1.45 V.s/cm^2^.

### LC-MS data analysis

The diaPASEF raw file processing and controlling peptide and protein level false discovery rates, assembling proteins from peptides, and protein quantification from peptides were performed using library free analysis in DIA-NN 1.8.^72^ Library free mode performs an in silico digestion of a given protein sequence database alongside deep learning-based predictions to extract the DIA precursor data into a collection of MS2 spectra. The search results are then used to generate a spectral library which is then employed for the targeted analysis of the DIA data searched against a Swissprot human database (January 2021). Database search criteria largely followed the default settings for directDIA including: tryptic with two missed cleavages, carbamidomethylation of cysteine, and oxidation of methionine and precursor Q-value (FDR) cut-off of 0.01. Precursor quantification strategy was set to Robust LC (high accuracy) with RT-dependent cross run normalization. Proteins with low sum of abundance (<2,000 x no. of treatments) were excluded from further analysis and resulting data was filtered to only include proteins that had a minimum of 3 counts in at least 4 replicates of each independent comparison of treatment sample to the DMSO control. Protein with missing values were imputed by random selection from a Gaussian distribution either with a mean of the non-missing values for that treatment group or with a mean equal to the median of the background (in cases when all values for a treatment group are missing) using in-house scripts in the R framework (R Development Core Team, 2014). Significant changes comparing the relative protein abundance of these treatment to DMSO control comparisons were assessed by moderated t test as implemented in the limma package within the R framework^73^.

### Co-immunoprecipitation

HEK293T cells were seeded into 6-well plate (3 × 10^5^ cells/well), cultured overnight, and then transfected with 1.5 μg FLAG-tagged BRD4 plasmid using TransIT-LT1 transfection reagents (Mirus Bio, Madison, WI, USA). The transfected cells were cultured for another 36 h, pretreated with 1 μM bortezomib, and co-treated with either compound or DMSO for 4 h before collection. The cells were collected and lysed in Pierce IP Lysis Buffer (Thermo Fisher Scientific) with cOmplete Mini Protease Inhibitor Cocktail (Roche, Basel, Switzerland) for 30 min on ice and centrifuged for 30 min at 4 °C to remove the insoluble fraction. For immunoprecipitation, 20 μL of pre-cleaned anti-FLAG M2 magnetic beads (Sigma-Aldrich) were added to the lysates. The beads–lysate mix was incubated at 4 °C for overnight on a rotator. Beads were magnetically removed and washed three times with PBS, and the FLAG-Tagged protein was competitively eluted using 3× FLAG Peptide (ApexBio Technology, Houston, TX, USA). Immunoblotting was carried out as previously described.

### Construction of plasmids

A gBlock (IDT) containing a codon-optimized YPEL5-Gly-Ser-FLAG coding sequence was cloned by Gibson assembly into a lentiviral transfer vector (pN106, gift from S. Gourisankar). The construct features the EF-1alpha promoter to drive transgene expression and the mouse Pgk-1 promoter to drive the expression of the Blasticidin resistance gene. YPEL5 mutations were introduced by overlap extension PCR followed by reassembly with the original cut vector. sgRNA constructs were assembled according to standard procedures as described for BsmBI-cut pXPR_023 (gift from S. Corsello; see Addgene #202447 for example). The destination lentiviral transfer vector contains an sgRNA cloning site (U6 promoter) and Cas9-FLAG-P2A-PuroR (EF-1alpha promoter). Codon-optimized YPEL5 and sgRNA sequences are provided below.

>YPEL5_gs_FLAG_opt

ATGGGAAGGATCTTTTTGGATCATATTGGAGGGACACGGCTGTTCTCTTGTGCTAACTGCGATACAATCTTGACAAACAGGTCCGAACTGATCTCTACGAGGTTTACGGGAGCAACCGGCAGAGCGTTCTTGTTTAACAAGGTCGTAAATCTCCAGTATTCCGAGGTACAGGATCGCGTCATGCTCACCGGGAGACACATGGTCAGGGACGTGTCCTGCAAGAACTGTAATTCCAAACTCGGCTGGATTTATGAATTCGCAACAGAGGATTCACAAAGATATAAAGAAGGGCGCGTCATTTTGGAACGAGCACTTGTGCGAGAATCCGAGGGTTTTGAAGAGCACGTCCCGTCTGATAACTCAGGTAGCGATTACAAGGACGACGATGACAAGtaa

>sgNEG

caccgGGTGTGCGTATGAAGCAGTG

>WDR26_sg_1_f

caccgCAGCCTGAATGTCAATAACG

>WDR26_sg_2_f

caccgGAGAGTCTGTAAACGCCGTG

>WDR26_sg_3_f

caccgCCTCTTACCACAATAGCATG

>WDR26_sg_4_f

caccgGACATCCTGACTCTTGCATG

>YPEL5_sg_1_f

caccgAGTACAGTGAAGTTCAAGAT

>YPEL5_sg_2_f

caccgAGTGGCGCCTGTGAAACGAG

>YPEL5_sg_3_f

caccgGTTTGCACAAGAAAACAGAC

### Generation of knockout cells

HEK293T cells were transfected with either pXPR_023/WDR26 or pXPR_023/YPEL5, along with pMD2.G (Addgene #12259) and psPAX2 (Addgene #12260), using TransIT-LT1 transfection reagents (Mirus Bio) to produce lentivirus. After 48 h of transfection, the supernatant was collected, filtered, and added to HiBiT-BRD4 Jurkat cells in the presence of 5 µg/mL polybrene (APExBIO, cat. no. K2701). Following 48 h of infection, the infected cells were passaged and cultured in medium containing 2 µg/mL puromycin (Gibco, cat. no. A11138-03) for three days. The surviving cells were grown for an additional five days. The knockout was confirmed by immunoblotting using either anti-WDR26 or anti-YPEL5 antibody.

### Generation of YPEL5 mutant cells

HEK293T cells were transfected with either pN106/YPEL5-FLAG-WT, pN106/YPEL5-FLAG-T36A, pN106/YPEL5-FLAG-R41A, or pN106/YPEL5-FLAG-L62A, along with pMD2.G (Addgene #12259) and psPAX2 (Addgene #12260), using TransIT-LT1 transfection reagents (Mirus Bio) to produce lentivirus. After 48 h of transfection, the supernatant was collected, filtered, and added to YPEL5^-/-^ HiBiT-BRD4 Jurkat cells in the presence of 5 µg/mL polybrene (APExBIO, cat. no. K2701). Following 48 h of infection, the infected cells were passaged and cultured in medium containing 10 µg/mL blasticidin (Gibco, cat. no. A11139-03) for five days. The surviving cells were grown for an additional 7 days. The re-expression of YPEL5 was confirmed by immunoblotting using anti-FLAG antibody.

### Cloning and recombinant protein preparation

The cDNA of full-length BRD4 was a gift from Peter Howley (Addgene #14447; http://n2t.net/addgene:14447; RRID:Addgene_14447),^74^ whereas those of BRD2 and BRD3 were acquired from an in-house human cDNA library (Max Planck Institute of Biochemistry). All new constructs were generated with the Gibson assembly method^75^ and mutagenized using the QuikChange site-directed mutagenesis protocol (Agilent, Santa Clara, CA, USA). To prepare multigene DNA constructs for insect cell expression, multiple inserts encoding the GID/CTLH E3 subunits were combined with the biGBac assembly method^76^ into single baculoviral expression vectors. All plasmids used in this study are listed in the Extended Data Table 1.

#### Insect cell expression and purification

All GID/CTLH E3 complexes as well as the E1 enzyme UBA1 were expressed in *Trichoplusia ni* High Five insect cells (Thermo Fisher Scientific) following three rounds of baculovirus amplification in *Spodoptera frugiperda* Sf9 cells (Thermo Fisher Scientific). The infected High Five cells were grown in EX-CELL 420 Serum-Free Medium at 27 °C for 72 h before harvesting by centrifugation (15 min, 450xg) and resuspension of the pelleted cells in the cold lysis buffer containing 50 mM Tris pH 8, 150 mM NaCl, protease inhibitors (10 μg/ml leupeptin, 20 μg/ml aprotinin, EDTA-free Complete protease inhibitor tablet (Roche, 1 tablet per 50 mL of buffer), 1 mM PMSF) and 5 mM DTT. The resuspended cells were disrupted by sonication and centrifuged (30 min, 20,000xg) to remove cell debris.

All versions of the GID/CTLH E3 assemblies harbored a Twin-Strep tag fused to the TWA1 C-terminus^39^ and were affinity purified from insect cell lysates by Strep-Tactin affinity (IBA-Lifesciences, Göttingen, Germany). The eluted proteins were subjected to size exclusion chromatography (SEC) with a Superose 6 Increase 10/300 GL column (Cytiva, Marlborough, MA, USA) in the final buffer containing 25 mM HEPES pH 7.5, 150 mM NaCl and 1 mM DTT (buffer A, for biochemical assays) or 5 mM DTT (buffer B, for cryo-EM). The WDR26^ΔCTLH^ module (alone and in complex with YPEL5) was expressed as N-terminal GST fusion. Proteins were purified by glutathione affinity chromatography (Cytiva) and incubated with Tobacco Etch Virus (TEV) protease (4 °C, overnight) to liberate the GST tag. The digested samples were subjected to SEC with a Superdex 200 10/300 GL column (Cytiva).

#### Bacterial expression and purification

Substrates (the BET family proteins and NMNAT1) and other reagents for biochemical assays (GID4 substrate receptor, E2 enzyme UBE2H, ubiquitin) were expressed in the codon-enhanced *Escherichia coli* BL21 (DE3) RIL cells. Bacterial cells were transformed with respective plasmids (Extended Data Table 1) and grown in the Terrific Broth (TB) medium supplemented with appropriate antibiotics at 37 °C until optical density (OD) of 0.6. After lowering the temperature to 18 °C, protein expression was induced with 0.4 mM isopropyl β-D-thiogalactopyranoside (IPTG) and carried out for 18 h. Cells were collected by centrifugation (15 min, 4,500xg) and resuspended in the cold lysis buffer containing 50 mM Tris pH 8, 150 mM NaCl, 1 mM PMSF and 5 mM DTT.

All versions of BRD2, BRD3, BRD4 as well as UBE2H were expressed as N-terminal GST fusions and purified analogously to the insect cell-expressed GST-TEV-WDR26^ΔCTLH^ (described above). To fully phosphorylate the C-terminal extension of UBE2H for ubiquitylation assays (which boosts its GID/CTLH E3-dependent activity), it was co-expressed with the catalytic subunit of the kinase CK2 (CK2α-6xHis),^38^ and subjected to anion exchange chromatography with the HiTrap Q HP column (Cytiva) prior to final SEC in buffer A.

The 6xHis tag was appended to, respectively, N- or C-termini of GID4 (Δ1-99) and NMNAT1. After resuspending bacterial cells in the lysis buffer supplemented with 10 mM imidazole, proteins were captured with the Ni-NTA affinity chromatography (Sigma-Aldrich) and subjected to SEC in buffer A.

WT ubiquitin was expressed in a tagless form and purified with the glacial acetic acid method followed by gravity cation exchange chromatography (GE Healthcare) and SEC as described previously.^77^

### Fluorescent labeling of substrates for biochemical assays

All substrates were fluorescently labeled by sortase A-mediated fusion^78^ of fluorescein (FAM) or TAMRA-containing peptides to the proteins’ N-terminal Gly exposed upon co-translational cleavage of initiator Met (for GG-NMNAT1-6xHis) or TEV-mediated removal of the GST tag (for GG-GSGS-BRD2/BRD3/BRD4). Labeling reactions were performed for 30 minutes at room temperature by mixing 50 μM substrate, 250 μM fluorescent peptide (fluorophore-GSGG-LPETGG, synthesized in the MPIB Bioorganic Chemistry & Biophysics Core Facility) and 10 μM sortase A-6xHis in the reaction buffer containing 50 mM Tris-HCl pH 8, 150 mM NaCl and 10 mM CaCl_2_. For labeling the BET proteins, the labeling mixture was supplemented with 5 mM imidazole after the reaction and passed back over Ni-NTA resin to remove 6xHis-tagged sortase A. The fluorescent substrates were purified by SEC in buffer A.

### Native PAGE band shift assay

For initial qualitative analysis of ZZ1-induced ternary complex formation (Fig. 1f), 1 μM FAM-labeled BRD4_BD1_ was incubated with 3 μM WDR26^ΔCTLH^ or YPEL5-WDR26^ΔCTLH^ in the presence of 10 μM JQ1 (Sigma-Aldrich) or ZZ1 (at 2% final DMSO concentration) for 1 h in buffer A. Samples were mixed with a non-denaturing PAGE loading buffer containing 5% (v/v) glycerol, bromophenol blue and Tris-borate (TB) buffer (100 mM Tris, 100 mM boric acid) and run on a native PAGE gel (pre-run at 200 V for 5 minutes) for 50 minutes (130 V, 4 °C) in the TB buffer. Ternary complex formation was assessed by monitoring the downward shift of FAM-BRD4_BD1_ (detected in a fluorescent scan) towards the faster-migrating band corresponding to YPEL5-WDR26^ΔCTLH^ (visualized by Coomassie staining).

The native gel was freshly poured and contained 4.5% (w/v) acrylamide/bis-acrylamide 29:1, 2% (v/v) glycerol, TB buffer, TEMED (75 μL/100 mL gel solution) and 0.04% (w/v) APS.

### *In vitro* ubiquitylation assays

All ubiquitylation assays were performed at room temperature in the final reaction buffer containing 25 mM HEPES pH 7.5, 150 mM NaCl, 0.25 mM DTT, 5 mM ATP and 10 mM MgCl_2_. Samples at indicated time points were quenched by mixing an aliquot of the total reaction mixture with a reducing Laemmli buffer and subjected to SDS-PAGE. Substrate ubiquitylation was visualized by a fluorescent scan of SDS-PAGE gels using the Amersham Typhoon Imager 600 (GE Healthcare, Chicago, IL, USA).

The initial ubiquitylation assays identifying the ZZ1-co-opted GID/CTLH E3 catalytic assembly and the targeted BRD4 construct (Fig. 1e, Extended Data Fig. 2c, 2h) were set up by first mixing 0.5 μM GID/CTLH E3 (catalytic core or its supramolecular WDR26-dependent version assembled with different substrate receptor modules, summarized in Extended Data Fig. 2b), 0.25 μM fluorescent substrate (FAM-labeled BRD4 or TAMRA-labeled NMNAT1), 1 μM phosphorylated UBE2H and 10 μM JQ1 or ZZ1 (at 2% final DMSO concentration). The mixture was incubated at room temperature for 1 h and supplemented with 0.2 μM E1 UBA1 and 20 μM ubiquitin to initiate the reaction. To probe the effect of the N-degron substrate receptor GID4 (Extended Data Fig. 2c, 2h), 1 μM GID4 (Δ1-99) was included in the reaction mixtures containing the GID/CTLH E3s assembled with the GID4-adapter subunit ARMC8.

Having identified YPEL5 as the ZZ1-induced neosubstrate receptor, all the following assays were performed with the YPEL5-GID/CTLH E3 (the assembly of RANBP9-TWA1-ARMC8-RMND5A-MAEA-WDR26-YPEL5) at the analogous reaction conditions.

To ensure an equal level of complex subunits in the assay querying structure-based YPEL5 mutants (Fig. 4c), the YPEL5-GID/CTLH E3 complexes were expressed by co-infection of High Five cells with two baculoviruses encoding: 1) WDR26-GID/CTLH E3 assembly (RANBP9-TWA1-RMND5A-MAEA-WDR26) and 2) WT or mutant YPEL5. The identity of the co-expressed YPEL5 mutants was confirmed by intact mass analysis, whereas their similar level was assessed by visual inspection of Coomassie-stained SDS-PAGE gels after ubiquitylation reaction.

### Intact mass spectrometry analyses probing the ZZ1 mode-of-action

To analyze whether ZZ1’s mode-of-action involves formation of a covalent adduct (Extended Data Fig. 4b), we incubated 5 μM YPEL5-WDR26^ΔCTLH^ with 10 μM BRD4_BD1_ and 20 μM ZZ1 (at 2% final DMSO concentration) in buffer A at room temperature overnight and analyzed the samples by intact mass spectrometry (performed in the MPIB Mass Spectrometry Core Facility).

To monitor the DTT-triggered conversion of degraders to their sulfinic acid versions (Fig. 3b and Extended Data Fig. 6c), DMSO stocks of ZZ1 or ZZ2 were incubated with buffer B (containing 5 mM DTT) for 2 h at room temperature and analyzed by intact mass spectrometry along with their DMSO-diluted controls.

### Fluorescence polarization (FP) assays

#### Quantification of ternary complex formation

To quantify ternary complex formation (Fig. 3d, 5e), we employed fluorescence polarization (FP) assay monitoring association of the fluorescent FAM-BRD_BD1_ tracer with YPEL5-WDR26^ΔCTLH^ upon degrader titration. To set up binding reactions, the mixtures of 20 nM FAM-BRD_BD1_ and 100 nM YPEL5-WDR26^ΔCTLH^ or WDR26^ΔCTLH^ in the binding buffer (25 mM Hepes pH 7.5, 150 NaCl, 5 mM DTT, 1 mg/mL BSA and 0.1 % Tween 20) were combined with the equal volume of 2-fold dilution series of the degraders or JQ1 (prepared in DMSO and then diluted with the binding buffer to reach the final DMSO concentration of 2% for all titration points) and incubated for 1 h at room temperature before transferring to 384-well flat bottom black plates (Greiner). Fluorescence polarization was calculated by measuring perpendicular and parallel fluorescence intensity values using the excitation and emission wavelengths of 482 nm and 530 nm, respectively, in the CLARIOstar microplate reader (BMG LABTECH). Polarization values obtained from three independent experiments were plotted in PRISM v10.3.1 (GraphPad). The potency of the degraders was expressed as half maximal effective concentration (EC_50_) estimated by non-linear regression using the four-parameter [agonist] vs. response model. Due to the substantially enhanced reactivity of ZZ2, its sulfinic acid version was prepared by 1 h incubation with the binding buffer before mixing with the protein mix and analyzed by intact mass spectrometry to confirm the conversion (Extended Data Fig. 6d).

#### Real-time FP monitoring the rate of ternary complex formation

To probe relative rates of ternary complex formation induced by the non-converted and sulfinic acid versions of degraders (Extended Data Fig. 5c, Fig. 5f), fluorescence polarization was monitored upon combination of the mixture of 10 nM FAM-BRD_BD1_ and 50 nM YPEL5-WDR26^ΔCTLH^ in the binding buffer (containing 1 mM DTT) with 10 μM of DMSO-dissolved ZZ1, ZZ2 and their sulfinic acid derivatives (resulting in the final 2% DMSO concentration). Contact of non-converted degraders with the DTT-containing buffer initiated their activation, resulting in a gradual increase of polarization. Experiments were also performed with ZZ1 and ZZ2 pre-incubated with the binding buffer before adding them to the protein mixture. An analogous FP assay was performed to query the impact of DTT concentration on the rate of ternary complex formation (Extended Data Fig. 5e). Here, the concentration of DTT in the binding buffer used for preparing the tracer-E3 ligase mixture varied from 0.1 to 5 mM.

#### Competitive FP probing cooperative E3 ligase-neosubstrate interactions

To test whether the tight ternary complex is driven by cooperative degrader-induced E3 ligase-target interactions (Fig. 4e), we monitored propensities of ZZ1-SO_2_H and JQ1 to displace the BRD4_BD1_-bound JQ1-FITC tracer (Tocris Bioscience, Bristol, UK) in the presence and absence of YPEL5-WDR26^ΔCTLH^. To determine conditions for the competitive FP assay, we first performed the binary binding experiment in a non-competitive format (Extended Data Fig. 6b). A 2-fold dilution series of BRD4_BD1_ prepared in the binding buffer was mixed with equal volumes of 20 nM JQ1-FITC. Polarization data from three independent experiments were fit to the one site-binding model in PRISM v10.3.1 (GraphPad) to determine the equilibrium dissociation constant (K_D_) for JQ1-BRD_BD1_ interactions.

To ensure sufficient signal for the competitive assay, we identified BRD4_BD1_ concentration at which the FP signal reached ∼60% saturation (i.e. 30 nM). Subsequently, a 2-fold dilution series of unlabeled competitors ZZ1-SO_2_H or JQ1 (prepared in DMSO and then diluted with the binding buffer to reach the final DMSO concentration of 2% for all titration points) was combined with equal volumes of the mixture of 20 nM JQ1-FITC and 60 nM BRD4_BD1_ alone or pre-mixed with 600 nM YPEL5-WDR26^ΔCTLH^. This led to displacement of JQ-FITC from BRD4_BD1_, thus decreasing fluorescence polarization. The values of polarization from three independent experiments were plotted in PRISM v10.3.1 (GraphPad) using the four-parameter [inhibitor] vs. response model to estimate the half maximal inhibitory concentration (IC_50_), reflecting relative binding strengths of the titrated competitors towards BRD4_BD1_. The extent of cooperativity was expressed as the apparent cooperativity factor (α_app_) defined as a ratio between the IC_50_ values in the absence and presence of the harnessed E3 ligase.

### Single-particle cryo-EM

#### Sample preparation and imaging

The YPEL5-GID/CTLH E3 for structural studies was expressed by co-infection of High Five cells with three separate baculoviruses encoding: 1) the GID/CTLH E3 catalytic core (RANBP9-TWA1-RMND5A-MAEA), 2) WDR26 and 3) YPEL5). The peak fractions (excluding those partially overlapping with the void volume peak) of the SEC-purified complex were pooled and concentrated to 6 mg/mL. The cryo-EM sample was prepared by combining 2.7 μM (2.5 mg/mL) YPEL5-GID/CTLH E3, 11 μM BRD4_BD1_, and 20 μM ZZ1 or ZZ2 in buffer B (containing 5 mM DTT) and incubating the mixture for 1 h on ice. Shortly before plunging, samples were supplemented with the 0.01% octyl-β-glucoside (β-OG) detergent (Sigma-Aldrich),^43^ which mitigates complex disassembly/protein aggregation during plunging and facilitates particle distribution.

Cryo-EM grids were prepared using Vitrobot Mark IV (Thermo Fisher Scientific) operated at 4 °C and 100% humidity. 3.5 μl of samples were applied to glow-discharged holey carbon R1.2/1.3, Cu 300 mesh grids (Quantifoil, Jena, Germany), blotted with Whatman no. 1 filter paper (blot time: 3 s, blot force: 3) and vitrified by plunging into liquid ethane.

#### Data collection and processing

Details of cryo-EM data collection are listed in Extended Data Table 1, whereas the flowchart of data processing workflow is presented in Extended Data Fig. 3 and 7.

Grids were first pre-screened on either a Talos Arctica or Glacios transmission electron microscope (Thermo Fisher Scientific) operated at 200 kV, equipped with a Falcon III (Thermo Fisher Scientific) or K2 (Gatan, Pleasanton, CA, USA) direct electron detector, respectively. The high-resolution datasets were acquired on a Titan Krios microscope (Thermo Fisher Scientific) operated at 300 kV, equipped with a post-column GIF and a K3 Summit direct electron detector (Gatan) operating in a counting mode. The automated data acquisition was carried out with SerialEM,^79^ while the Focus software^80^ was used for the on-the-fly discarding of poor-quality images and data pre-processing.

Movie frames were motion-corrected with dose-weighting using MotionCor2^81^ and subjected to estimation of the contrast transfer function (CTF) with Gctf^82^ integrated in Focus. Particles were automatically picked with Gautomatch (K. Zhang) using our previously published map of the YPEL5-GID/CTLH E3 (EMDB: EMD-18170)^43^ as a template. All the subsequent stages of data processing were carried out with RELION 4.0 and 5.0.^83,84^ To clean up the data, while preserving particle views corresponding to less populated orientations, the extracted particles were subjected directly to three rounds of unmasked 3D classification. The final classification revealed the co-existence of multiple types of BRD4_BD1_-bound YPEL5-GID/CTLH E3 assemblies with similar shapes and dimensions but differing stoichiometries of the catalytic and YPEL5-WDR26 modules as previously reported.^43^ The compositional heterogeneity as well as the dynamic nature of the supramolecular GID/CTLH E3s limit the resolution of their overall maps.^38^ To gain molecular insights into the architecture of the ternary complex, we generated masks around the BRD4_BD1_-YPEL5-WDR26-containing regions in the 3D refined maps of each type of the GID/CTLH E3 assembly and extracted the encompassed densities by signal subtraction. In doing so, we not only overcame the challenge of complex heterogeneity but also substantially increased the number of particles available for subsequent 3D classifications. The generated pools of signal-subtracted particles were joined, 3D classified, and aligned by 3D refinement with a mask over the BRD4_BD1_-bound YPEL5-WDR26 and the better-resolved copy of the scaffolding module (RANBP9-TWA1). The map was further improved by local refinement with a mask around the BRD4_BD1_-bound YPEL5-WDR26 module. Additional rounds of focused 3D classification (without particle alignment) enriching for particles with the most complete and well-resolved features followed by local refinements with Blush regularization^85^ yielded the final high-resolution reconstructions permitting model building. For the map of the ZZ2-SO_2_H-induced ternary complex, the final round of local refinement was preceded by CTF refinement. Maps were post-processed by B-factor sharpening and high-resolution noise substitution in RELION. To aid in building the atomic models, the refined maps were also sharpened with DeepEMhancer^67^ (employing the ‘highRes’ model) and are deposited as additional maps in EMDB. The estimated resolutions of final maps are based on the gold-standard Fourier Shell Correlation (FSC) at a 0.143 cut-off.

#### Model building, refinement and analysis

Atomic models were manually built and refined with Coot v0.9.8.7,^86^ whereas degrader coordinates and the corresponding restraints applied during model refinement were generated using the JLigand^87^ interface. The refinement and validation statistics of the built models are listed in Extended Data Table 1.

The structure of the ZZ1-SO_2_H-induced ternary complex was obtained by first docking the previous coordinates of the YPEL5-WDR26 module (PDB: 8QBN)^43^ and BRD4_BD1_ (PDB: 3MXF)^34^ in UCSF Chimera^88^ v1.11.2 into the post-processed map and manually refining the differing parts (e.g. removing the parts of the BRD4 bromodomain facing away from the YPEL5 interface, which are less ordered and thus blurry/invisible in the high-resolution reconstruction). This left an unoccupied segment of electron density filling the canonical acetyl-lysine binding pocket of BRD4_BD1_ and the central basic groove of YPEL5 that supported fitting ZZ1-SO_2_H. The coordinates of the -SO_2_H group (wherein the sulfur center adopts the pyramidal geometry and features the lone pair of electrons) were oriented to best fit the map while maximizing polar interactions with the surrounding YPEL5 sidechains. Prominent blobs of electron density at the three-way interface of ZZ1-SO_2_H chloro, YPEL5 and BRD4_BD1_ were assigned as water molecules, which appear to participate in formation of a cooperative ternary complex. To obtain the structure of the ternary complex with the improved degrader ZZ2-SO_2_H, we docked the coordinates of the ZZ1-SO_2_H-driven complex and manually refined differing areas. As in the first structure, the degrader density was clearly resolved (including the region corresponding to the appended chloro group) and enabled its unambiguous fitting.

Both models were subjected to iterative rounds of manual building and real-space refinement in Phenix v1.21.1^89^ until a satisfactory model quality, in terms of geometry and agreement with the map, was achieved. Configurations of the zinc-binding sites within WDR26 and YPEL5 as well as that of the degraders were restrained during real-space refinement. The final model was validated with Molprobity.^90^ Structure visualization and analyses were carried out with UCSF Chimera v1.11.2^88^, UCSF ChimeraX v1.7.1^91^ and PyMOL v3.0.3 (Schrödinger, New York, NY, USA). The surface area of the YPEL5-BRD4_BD1_ interface in the ternary complex structure was calculated with the PISA v1.52 tool of the European Bioinformatics Institute.^47^

### LC-MS detection of ZZ1

Following treatment of cells with 5 μM of ZZ1 for 5 h, cells were washed 2 times with PBS and lysed in 100 ul of a 2:1:1 mixture of acetonitrile: methanol: water. The cell pellets were vortexed well to ensure complete lysis, centrifuged at 4 °C for 10 min at 15,000 rpm and the supernatant was transferred to a LC-MS vial.

Untargeted metabolomics measurements were performed using an Agilent 6530 Quadrupole time-of-flight LC-MS instrument. MS analysis was performed using electrospray ionization (ESI) in positive mode. The dual ESI source parameters were set as follows: the gas temperature at 250 °C; the drying gas flow at 8 l/min; the nebulizer pressure at 25 psi; the sheath gas temp at 300 °C; the sheath gas flow at 12 l/min; the capillary voltage at 3500 V; and the fragmentor voltage at 175 V. Separation of metabolites was conducted using an Eclipse Plus C18 LC column (Agilent, cat. no. 959961-902). Mobile phases were as follows: buffer A, water with 0.1% formic acid; buffer B, acetonitrile with 0.1% formic acid. The LC gradient started at 70% B and increased to 100% B from 0-4 min then held constant at 100% B from 4-5 min before going back down to 70% B from 5-5.1 min. The flow rate was at a constant 0.7 ml/min through the entire run.

## Acknowledgements

This work was support by the National Institutes of Health (NIH) grants P01CA066996 and R35CA253125 (to B.L.E.), R01CA262188 and R01CA2144608 (to E.S.F.), R01CA218278 (to N.S.G. and E.S.F.), NIH High End Instrumentation grant (1S10OD028697-01) (to N.S.G.), the Howard Hughes Medical Institute (to B.L.E.), and departmental funds from Stanford Chemical and Systems Biology and Stanford Cancer Institute (to N.S.G.). W.S.B. is supported by Basic Science Research Program through the National Research Foundation of Korea (NRF) funded by the Ministry of Education (grant no. RS-2024-00410290). Z.K. is supported by Swiss National Science Foundation (grant no. P500PB_214385). J.C., S.S., and B.A.S. were supported by the Max Planck Gesellschaft, the European Union (ERC, UPSmeetMet, 101098161), and Leibniz Prize from the Deutsche Forschungsgemeinschaft (DFG, SCHU 3196/1). We thank S. Übel and S. Pettera from MPIB Bioorganic Chemistry & Biophysics Core Facility for peptide synthesis; D. Bollschweiler and T. Schäfer for cryo-EM assistance; B. Steigenberger and V. Sanchez from the MPIB Mass Spectrometry Core Facility for intact mass analyses, J. Rajan Prabu for guidance in structural analysis; S. von Gronau, J. Kellermann and members of the Schulman lab for advice and support.

## Author contributions

Z.Z., W.S.B., and J.C. contributed equally to this work. Z.Z., W.S.B., J.C., B.A.S., and N.S.G. conceived and initiated the study; Z.Z. designed and synthesized molecular glue degraders; W.S.B. designed and conducted cell biological experiments with the help of S.M.H. and I.Y.; J.C. designed and conducted biochemical and structural biology experiments with the help of S.S.; Z.K. carried out Bison CRISPR screening with the help of M.S.; V.L.L. carried out HPLC-MS analysis; D.M.A. and K.A.D. performed whole-cell proteomics experiments; N.S.G., B.A.S, B.L.E., and E.S.F. supervised the project; The manuscript was written by Z.Z., W.S.B., J.C., Z.K., S.M.H., B.A.S, and N.S.G. with input from all authors.

## Competing interests

N.S.G. is a founder, science advisory board member (SAB) and equity holder in Syros, C4, Allorion, Lighthorse, Voronoi, Inception, Matchpoint, CobroVentures, GSK, Shenandoah (board member), Larkspur (board member) and Soltego (board member). The Gray lab receives or has received research funding from Novartis, Takeda, Astellas, Taiho, Jansen, Kinogen, Arbella, Deerfield, Springworks, Interline and Sanofi. B.A.S. is a member of the scientific advisory board of Proxygen, and co-inventor of intellectual property licensed to Cinsano. B.L.E. has received research funding from Novartis and Calico. He has received consulting fees from Abbvie. He is a member of the scientific advisory board and shareholder for Neomorph Inc., TenSixteen Bio, Skyhawk Therapeutics, and Exo Therapeutics. E.S.F. is a founder, scientific advisory board (SAB) member, and equity holder of Civetta Therapeutics, Proximity Therapeutics, and Neomorph, Inc. (also board of directors). He is an equity holder and SAB member for Avilar Therapeutics, Photys Therapeutics, and Ajax Therapeutics and an equity holder in Lighthorse Therapeutics. E.S.F. is a consultant to Novartis, EcoR1 capital, Odyssey and Deerfield. The Fischer lab receives or has received research funding from Deerfield, Novartis, Ajax, Interline, Bayer and Astellas. K.A.D. receives or has received consulting fees from Kronos Bio and Neomorph Inc. M.S. has received research funding from Calico Life Sciences LLC. All other authors declare no competing interests.

## Data availability

Data generated in this study are provided in the manuscript, supplementary information, and source data files. Additional data supporting the findings of this study are available from the corresponding author upon reasonable request. Cryo-EM density maps and atomic models have been deposited in the Electron Microscopy Data Bank (EMDB) and the RCSB Protein Data Bank (RCSB PDB), respectively, and will be released upon publication.

